# Dorsal CA1 Hippocampal Place Cells Form a Multi-Scale Representation of Megaspace

**DOI:** 10.1101/2021.02.15.431172

**Authors:** B.C. Harland, M. Contreras, M. Souder, J.M. Fellous

## Abstract

Spatially firing “place cells” within the hippocampal CA1 region form internal maps of the environment necessary for navigation and memory. In rodents, these neurons have been almost exclusively studied in small environments (<4 m^2^). It remains unclear how place cells encode a very large open 2D environment, which is more analogous to the natural environments experienced by rodents and other mammals. Such an ethologically realistic environment would require a more complex spatial representation, capable of simultaneously representing space at overlapping multiple fine to coarse informational scales. Here we show that in a ‘megaspace’ (18.6 m^2^), the majority of dorsal CA1 place cells exhibited multiple place subfields of different sizes, akin to those observed along the septo-temporal axis. Furthermore, the total area covered by the subfields of each cell was not correlated with the number of subfields, and this total area increased with the scale of the environment. The multiple different-sized subfields exhibited by place cells in the megaspace suggest that the ensemble population of subfields form a multi-scale representation of space within the dorsal hippocampus. Our findings point to a new dorsal hippocampus ensemble coding scheme that simultaneously supports navigational processes at both fine- and coarse-grained resolutions.

## Introduction

Seminal place cell studies found that the majority of place cells formed a single field in the ‘classic’ environments (<1 m^2^) tested [1–3]. When such environments were expanded, place field size also expanded [2, 4]. It was also shown that individual place cells along the dorso-ventral axis of the hippocampus coded areas of increasingly larger sizes [1, 5]. Taken together, these findings suggested that the larger ventral hippocampus place fields may be involved in representing large-scale environments. How this multi-scale information is effectively integrated and used is however unknown, especially given the dynamic nature of the code as observed through, for example, remapping experiments [6]. An alternative theory is that the multi-scale nature of the spatial code is not purely predicated on the anatomical location of place cells, and that it is the result of dynamic ensemble coding throughout the entirety of the hippocampus [7]. However, experimental support for ensemble place cell coding of multiple spatial scales has, to this day, been lacking.

Fenton et al. (2008) showed that in a larger classic environment (2.1 m^2^), place cells have multiple irregularly arranged, enlarged place fields. Since then, several studies have further reported multi-field place cells on long linear running tracks (10.3m, 18m, and 48m)[5, 8, 9] in rats, and a 200m tunnel in bats [10]. However, because animals are constrained to run in a particular direction in these linear environments, place cells operate differently than in open-fields, by forming for example, bi-directional selectivity [11]. In a large open-field arena (2.5 m^2^), Park et al. (2011) showed multiple field place cells in dorsal hippocampal CA1, CA3, and dentate gyrus. While the area of the largest subfield per cell increased on average from a small to a larger environment, no significant change in area was noted when all subfields were accounted for, unlike previous studies which showed that the average field size increased [2, 12]. Overall, this experimental work challenged existing place cell models, which were based on the idea of one-place-cell / one-location. This resulted in an alternative computational model positing the existence of a ‘megamap’ in which individual place cells feature multiple subfields of similar sizes, capable of enlarging to fill any infinite space [13]. Experimentally, it is still unclear how enlarged multiple-field place cells would effectively encode a large ‘megaspace’ at multiple spatial scales. Understanding how place cells encode multiple spatial scales has additional theoretical value as these same hippocampal neurons are thought to be involved in encoding human autobiographical memory along multi-scale mnemonic hierarchies [14–16].

Here, we compared place cell properties in a megaspace (18.6 m^2^), considerably larger than previous published studies, with those in a classic environment. We used wireless recording and a new behavioural paradigm in which rats were trained to follow a small food-baited robot to obtain place cell recordings with sufficient coverage within the megaspace. We found that place cells exhibited multiple spatially distributed subfields of many sizes in the megaspace. We found that the average place field size increased with the size of the environment. We also show that the subfields of each individual cell were of different sizes and that the number of subfields per cell was not correlated with the total area covered by a cells subfields.

## Results

### Robot following facilitated high resolution place cell recordings in the megaspace

Rats were recorded in a megaspace (5.3 x 3.5 m; 18.6 m^2^, Fig. 1A, Movie S1) in between visits to a smaller environment (1.8 x 1.2 m; 2.2 m^2^). The megaspace is considerably larger than environments used in previously published studies to record place cells (Fig. 1B). To obtain sufficient coverage of this environment, we trained rats to follow a small food-baited robot (‘Sphero’) controlled by an experimenter (Fig. 1C). Previously, we have shown that rats can attend to their surrounding by learning an allocentric spatial task while following the robot and that place cells did not remap in a small environment during robot following [17]. Here, we compared behavioural and place cell parameters between separate robot following (N = 39) and traditional foraging (N = 15; Fig 1D) sessions using one-way Anova (see Fig. S1A-C for example trajectories). In the megaspace, robot-following ensured greater behavioural coverage (*F*_(1,53)_ = 25.43, *P* < 0.0001; Fig. 1E) and movement velocity (*F*_(1,53)_ = 54.26, *P* < 0.0001; Fig. 1F) compared with foraging. Similarly, coverage and velocity were also increased in the small environment in robot-following sessions (Fig. S1D-E).

**Fig. 1:**
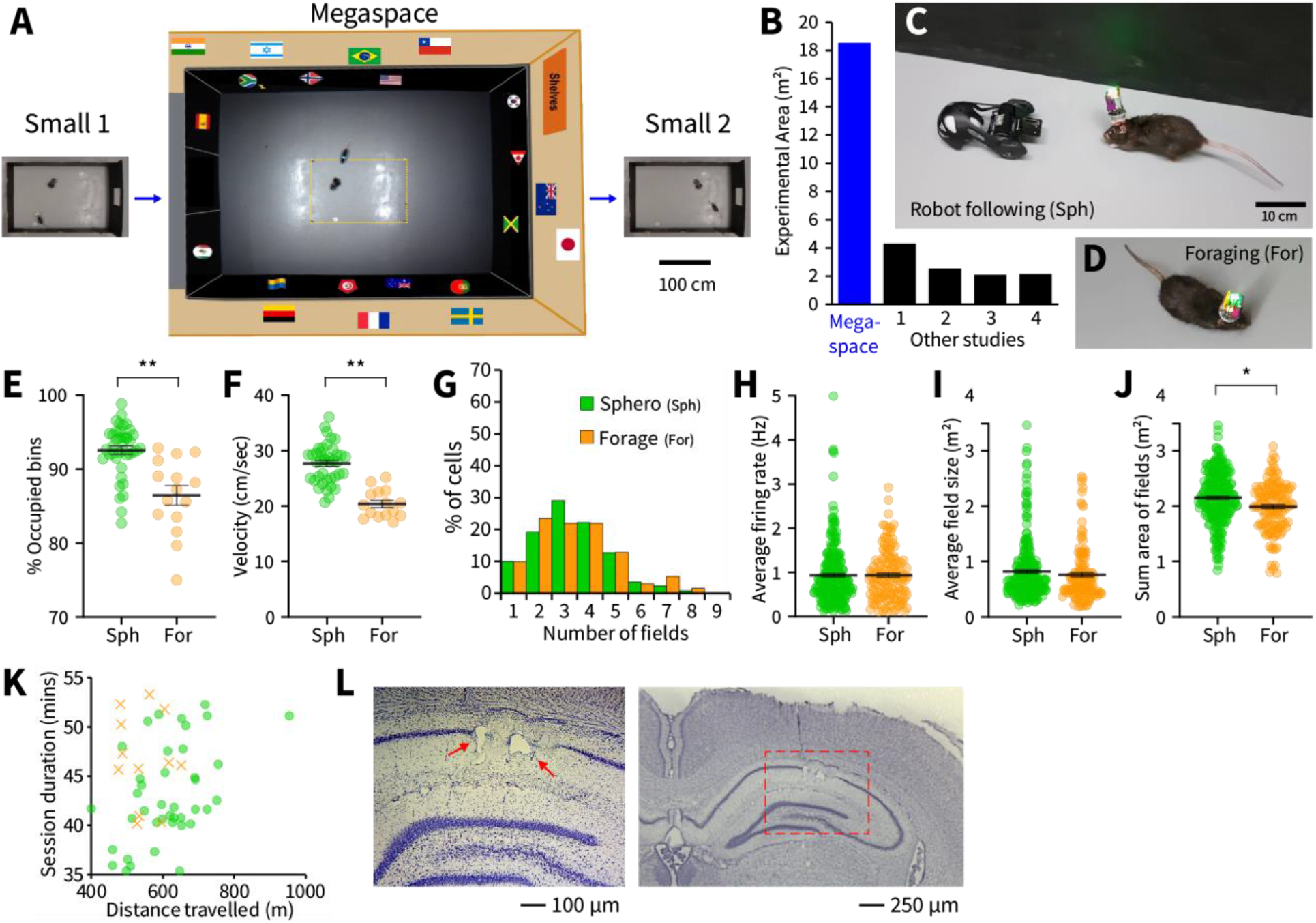
Methods and comparisons between robot-following and foraging sessions. (**A**) Top view of recording environments. Yellow dotted line shows position of small environment within the megaspace (18.6 m^2^). (**B**) The megaspace is over four times larger than environments from other published studies which also included dorsal CA1 place cell recordings: 1 = 48m track [9]; 2 = Large box [7]; 3 = Monkey cage [12]; 4 = 18m track [5]. (**C**) Rats were trained to follow a small baited robot (‘Sphero’). A wireless headstage allowed for recordings in the megaspace. Robot-following was compared with (**D**) traditional foraging. (**E**) Robot-following (Sph, green) resulted in a greater fraction of the room covered by the occupancy map, and (**F**) greater average speed in the megaspace than during classic foraging (For, orange). Place cells in the megaspace had similar (**G**) numbers of subfields, (**H**) average firing rates, and (**I**) average place field sizes in robot following and foraging sessions. (**J**) The sum area of place fields per cell was greater in robot following sessions in the megaspace. (**K**) Robot-following sessions (green circles) yielded more distance traveled in a smaller amount of time compared with foraging (orange crosses). (**L**) Coronal section showing dorsal hippocampus. Arrowheads show electrolytic lesions indicating the end of tetrode tracks. For all panels * = *P* <0.05, ** = *P* <0.001.

There were no significant differences in place cell characteristics in the small environment between robot following and foraging sessions (Fig. S1F-I). Place cell characteristics in the megaspace did not differ between session types for number of fields (*F*_(1,381)_ = 0.33, *P* = 0.57; Fig. 1G), average firing rate (*F*_(1,381)_ = 0.05, *P* = 0.83; Fig. 1H), and mean size of place fields (*F*_(1,381)_ = 1.15, *P* = 0.28; Fig. 1I). The total area of place subfields for a given cell in the megaspace was slightly higher for cells in robot-following sessions (mean=2.15m^2^; SD±0.47m^2^) than for cells during foraging (mean=1.99m^2^; SD=0.44m^2^; *F*_(1,381)_ = 10.57, *P* < 0.01; Fig. 1J). This difference was due to the lower average velocity in foraging sessions; when a sub-set of velocity-matched robot following and foraging sessions (n = 6 each) were compared, there were no differences in place cell characteristics, including total area of place subfields in the megaspace (Fig. S1L-P). Robot following generally resulted in greater distances travelled in a shorter time (Fig. 1K) without altering place cell function, therefore place cells in robot-following and foraging sessions were pooled for all further analyses.

### Most place cell had multiple subfields of different size in the megaspace

We recorded 539 place cells from dorsal CA1 over 54 sessions in five rats (Small 1 – Megaspace – Small 2; Fig. 1A). Tetrode positions in the dorsal CA1 were confirmed histologically (Fig. 1L) and the position of each tetrode analyzed was verified to be within the CA1 area of the hippocampus (Fig. S2). To ensure that activity in the megaspace could not be explained by tetrode drift over the long sessions, only spatially firing place cells active in all three environments, with stable place fields in both small environments, were retained for analysis (n = 383 place cells; 71% of total place cells). We compared place cell characteristics between small 1, small 2, and the megaspace using one-way Anova’s and Tukey’s HSD tests.

Most place cells had multiple subfields with a broad range of sizes in the megaspace (Fig. 2A; more examples shown in Fig. S3), exhibiting more spatial subfields per cell compared to within the small environments (*F*_(2,1146)_ = 405.6, *P*’s < 0.0001). The majority of cells (82%) exhibited 2-5 subfields in the megaspace compared to 1-2 subfields (91%) in the small environments (Fig. 2B). Place subfields in the megaspace were also significantly larger on average, both in terms of the mean area of their subfields (*F*_(2,1146)_ = 560.2, *P’s* < 0.0001; Fig. 2C) and their sum area (*F*_(2,1146)_ = 6203.5, *P’s* < 0.0001; Fig. 2D). The up-scaled multiple-subfield representation yielded only 2% more relative coverage per place cell in the megaspace than in the small environment (*F*_(2,1146)_ = 95.4, *P’s* < 0.0001; Fig. 2E), which was 8.8 times smaller in overall area. This coverage difference was reduced to 0.9% in a sample of megaspace and small environment visits matched for rat velocity (data not shown). Number of subfields, mean and sum area of subfields, and percentage of environment did not differ between Small 1 and Small 2 (Tukey’s post-hoc comparisons, *P’s* > 0.85). Cells exhibited a comparable average firing rate in the different sized environments (*F*_(2,1146)_ = 1.92, *P* = 0.15), and the average firing rate in the megaspace was 0.93 ± 0.64 SD (Fig. 2F).

**Fig. 2:**
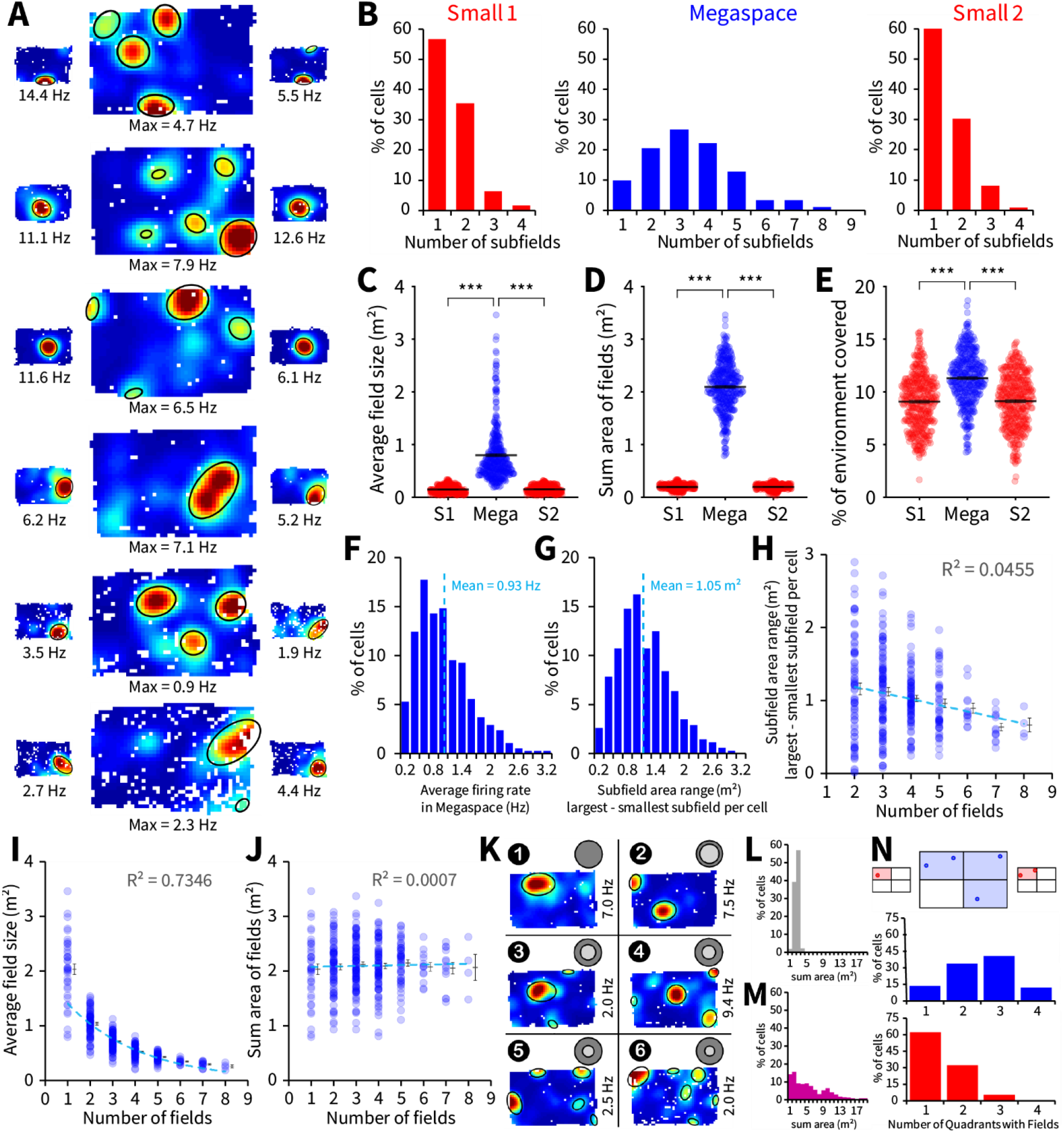
In the megaspace, place cells had multiple subfields of various sizes. (**A**) Six different representative place cells: top four cells recorded with robot-following, bottom two with foraging. Place cells exhibited multiple subfields of varying size in the megaspace. (**B**) Number of place subfields per cell for the three recording epochs. (**C**) Mean and (**D**) sum area of all subfields per cell was significantly greater in the megaspace although (**E**) only ~2% more space is covered compared to the small environment. (**F**) Average firing rate of place cells in the megaspace, only cells with > 0.1 Hz average firing rate were considered place cells. (**G**) Most cells with at least two subfields in the megaspace had a range of subfield sizes (area of largest – smallest subfield per cell) greater than 0.6m^2^. (**H**) A linear trend suggested that most place cells could possess subfield sizes of multiple scales, irrespective of their number of subfields. (**I**) In the megaspace, the average subfield size decreases with the number of subfields per cell, however (**J**) the subfield sum area of cells was not correlated with the number of subfields. (**K**) Example of place cells with 1 – 6 subfields, which have the mean sum area of fields shown in the outer dark-grey filled circle above each rate map, and the mean place field area shown in the inner light-grey filled circle. Number of subfields indicated above each graph, on the left. Sum area of subfields per cell is shown when (**L**) a cell-specific (> 1.2 SD above mean) and (**M**) fixed (> 1 Hz) firing rate threshold are used to define place fields, 45 out of 383 cells did not have fields using the fixed threshold. (**N**) Diagram showing a representative cell quantified for the number of quadrants containing subfields in each environment. The average number of quadrants occupied per cell with subfield centers is shown for the megaspace (blue) and small environment (red). For all panels *** = *P* <0.0001.

Most multi-field place cells in the megaspace had subfields that greatly ranged in size, with most cells (79%) having a subfield area range greater than 0.6 m^2^ (Fig. 2G). There was only a weak negative correlation between subfield area range and number of subfields suggesting cells with either few or many subfields had the capacity to be ‘multi-scale’ in the megaspace (Fig. 2H). We confirmed that place subfield locations were not correlated between the small environment and megaspace within each sessions (Fig. S4A-E), and that in a smaller ‘classic environment’ (<1m^2^), most place cells (94%) had only one place field (Fig. S4F-I, ‘very small’).

### In megaspace, the number of subfields a cell had was not correlated with the total area of floor space covered by the cell

We next investigated the relationship between the number of subfields per cell and the megaspace area covered by those subfields. We found that the more subfields a cell possessed, the smaller the average size of these subfields, as described by a negative exponential distribution (r = 0.86; Fig. 2I). In contrast, the average total area of all subfields remained constant irrespective of the numbers of subfields (linear fit, r = 0.027, Fig. 2J). Figure 2K, shows examples of different place cells with 1 to 6 subfields, each covering this average sum area of all subfields. However, comparing sum area across cells is problematic as the cell-specific threshold used for labelling place fields and the rejection of low firing rate cells (< 0.1 Hz) resulted in a narrow range of subfield sum areas (Fig. 2L, S4J) compared to the wide range of sum areas evident when a 1 Hz threshold to define place fields was applied across all cells (Fig. 2M, also see S5).

One advantage of having multiple subfields is to allow each cell to contribute to an ensemble spatial representation in multiple regions within the environment. Indeed, most place cells in the megaspace had subfields distributed in 2-4 quadrants (86%), with the highest proportion covering 3 (Fig. 2N). In contrast, place cell’s subfields in the small environment mostly covered 1-2 quadrants (95%) with the highest proportion of cells covering only 1.

The finding that cells in the megaspace possessed a wider range of place subfield sizes, persisted when we changed the bin-size (Fig. S4K-N), included a larger population of place cells (Fig. S4O-Q), used the fixed 1 Hz common threshold for labelling place fields (Fig. S5), and when a range of cell-specific firing rate thresholds for subfield detection were applied (Fig. S6). The lack of correlation between number of subfields and sum area of subfields also persisted but was slightly negatively correlated when the 1 Hz threshold was applied (Fig. S5E).

### The ensemble of subfields in the megaspace formed a multi-scale representation of space

A subset of 125 (out of 383) cells were used to plot the set of ‘idealized’ (circular, equivalent area) place fields in the three environments, categorized into seven color-coded size ranges (Fig. 3A). This subset included only one cell (best isolation) per tetrode per session, and was used to ensure that findings were not contaminated by overlap errors in cluster cutting. Place fields in the smaller environments were all of a similar scale, mostly falling into the two smallest color-bands (purple and dark blue, Fig. 3A). In contrast, fields in the megaspace were of many different scales forming a near-uniform multi-scale representation of space (Fig. 3B) with increased variability of subfield sizes (Fig. 3C). A major functional advantage of the multiple-place field representation in the megaspace was a significant increase in the number of overlapping subfields (One-way Anova: *F*_(1,198)_ = 12.21, *P* < 0.001; Fig. 3D) compared with the small environment. Such highly overlapping ensemble patterns of activity within a population of place cells can in principle accurately estimate location [7].

**Fig. 3:**
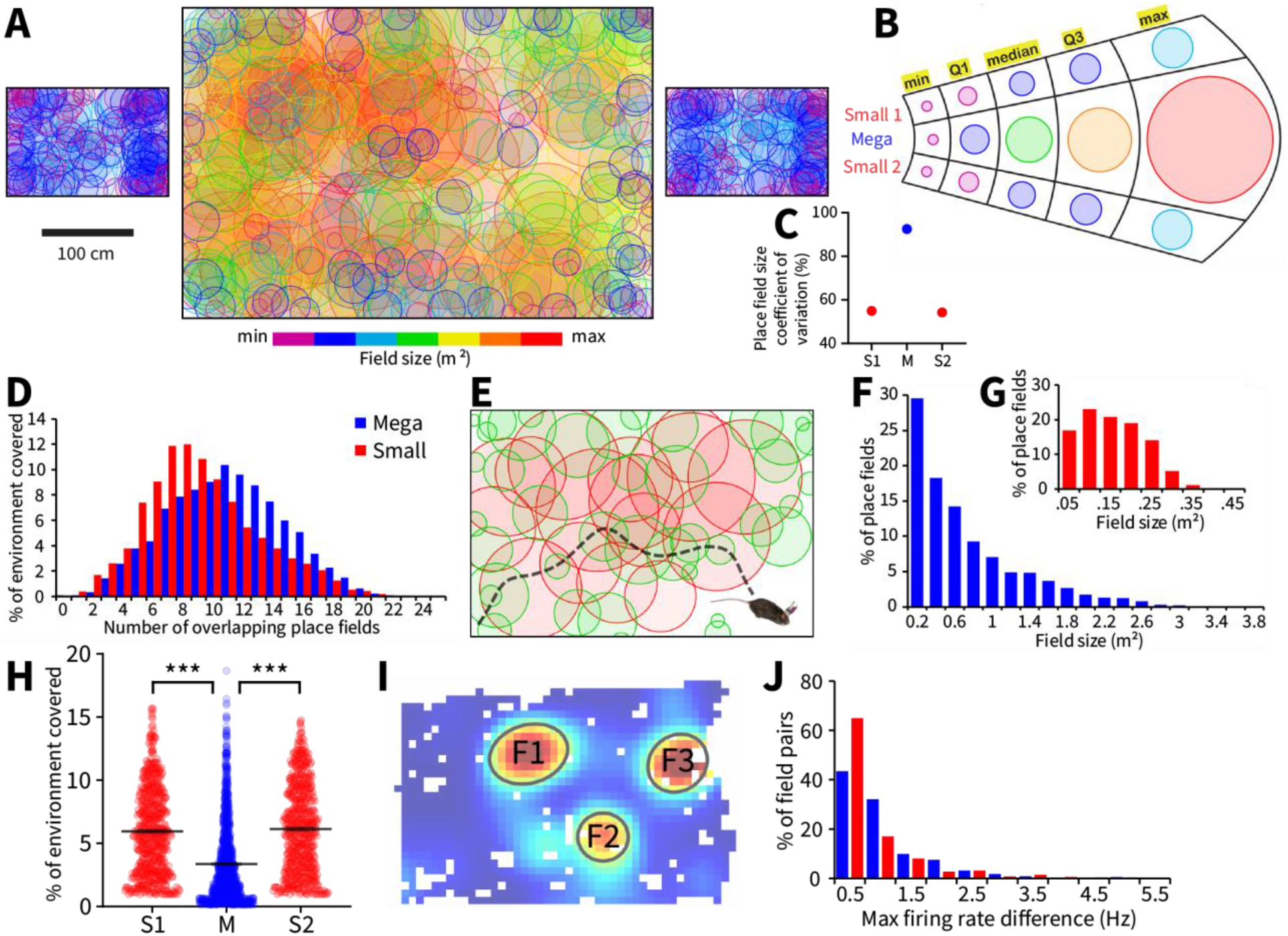
The population of place subfields formed a multi-scale representation of space in the megaspace. (**A**) Population of subfields from 125 well isolated place cells plotted in seven color-bands based on their area in the megaspace from smallest (purple, 0.023 – 0.091 m^2^) to largest (red, 1.23 – 3.46 m^2^). (**B**) There was a greater range of place field size in the megaspace than in the small environments, (**C**) reflected by the greater variability in field size (**D**). There was a greater degree of place subfield overlapping in the megaspace compared with the small environment. (**E**) Cartoon illustrating the prediction that many more smaller place subfields (green circles, n = 40) would be required in order to support finer-grain representations of the megaspace than large subfields (red circles, n = 16) would be needed to support coarser-grained representations. (**F**) The distribution of subfield sizes in the megaspace was consistent with this prediction (n = 1288 subfields). (**G**) There was a different distribution of subfield sizes in the small environment (Small 1 and Small 2; n = 1152 subfields). (**H**) % of environment covered per place field. (**I**) The difference between maximum firing rate of each pair of subfields was calculated, |F1-F2|, |F1-F3|, |F2-F3|. (**J**) Distributions of differences in maximum firing rate between subfield pairs from all cells are shown for the megaspace (blue, n = 1964 subfield pairs) and small environment (red, n = 468 subfield pairs). These differences are typically small, which suggests that subfield firing rate is not sufficient to differentiate spatial position for multiple subfield place cells. For all panels *** = *P* <0.0001.

This ensemble of small and large overlapping place fields may contribute to both fine- and coarse-grained spatial representations of the environment as well as to the disambiguation of spatial location, with each location within the megaspace being uniquely characterized by a specific set of subfields of different sizes. Coarse-grained representations would support fast traversal of open space at the scale of meters whereas fine-grained representations would support higher resolution navigational operations at the scale of centimeters. A coarse-grained spatial representation would consist of very large overlapping place fields and would require much fewer fields than a fine-grained representation of smaller overlapping place fields (Fig. 3E). This may explain the distribution of place field sizes in the megaspace (Fig. 3F) which is well fitted by a negative exponential curve (r = 0.995) with the majority of fields (78%) having an area of 1m^2^ or less, and the remainder (22%) between 1m^2^ and 4m^2^. In contrast, in the smaller environments, the distribution of subfield sizes was well fitted by a Gaussian function (r = 0.985; Fig. 3G), although it became quasi linear when lower thresholds for labelling subfields were applied (Fig. S6F). Individual place subfields covered less fraction of the environment in the megaspace compared to the small environment (One-way Anova: *F*_(2,2440)_ = 196.37, *P* < 0.0001; Fig. 3H). We tested whether each cell’s subfields had different peak firing rates, which could allow for within-cell differentiation of spatial position (Fig. 3I) and found that these firing rate differences were small, both in the megaspace and the small environments (Fig. 3J). Altogether, these data suggest that small and large overlapping place fields from many simultaneously active place cells form a multi-scale ensemble representation of the animal’s position within the megaspace.

### Place cells exhibited irregular patterns of subfields distribution across the megaspace

The distribution and spatial position of the population of place subfields were consistent with an ensemble coding scheme of spatial position in which the population discharges at each location are unique [7]. The population of subfield centers from all place cells was spread out within the megaspace, with no evidence of clusters or repeated positional patterns (Fig. 4A). There was a small accumulation of fields near the walls, possibly because walls were ‘cue-rich’, whereas fields located in the rooms center were ‘cue-poor’ [18]. However, there was only a moderate positive linear correlation (r = 0.36) between subfield size and distance to the closest wall in the megaspace (Fig. 4B). Interestingly, there was more wall clustering (Fig. 4C) and a stronger positive linear correlation (r = 0.40) between field size and distance to closest wall in the small environment than in the megaspace (Fig. 4D), despite the more limited range of subfield sizes. We quantified the percentage of place fields in each environment that contacted the walls, the corners, and those that did not contact any boundary, “middle cells” (Fig. 4E). In spite of the very different environmental scales, and characteristics of place cells in the different environment, there were similar proportions of corner, wall and middle located subfields in the megaspace and small environment (Fig. 4F). We next investigated the distance between subfields within each cell in the megaspace. The configurations of individual place fields per cell in the megaspace appeared to be irregular [12] as evidenced by the fact that they were normally distributed both for average distance between field centers, (Kolmogorov-Smirnov test; *D*_(345)_ = 0.044, *P* > 0.2; Fig. 4G) and field edges (*D*_(345)_ = 0.037, *P* > 0.2; Fig. 4H). Randomly generated place field positions in the megaspace (Fig. 4I) also produced a normally distributed pattern of average distances between place fields per cell (*D*_(345)_ = 0.057, *P* > 0.2), however, slightly offset to the left towards lower average distances (Fig. 4J). The more abundant fields in the corners in the data resulted in more high distance field pairs compared to the more dispersed simulated field centers.

**Fig. 4:**
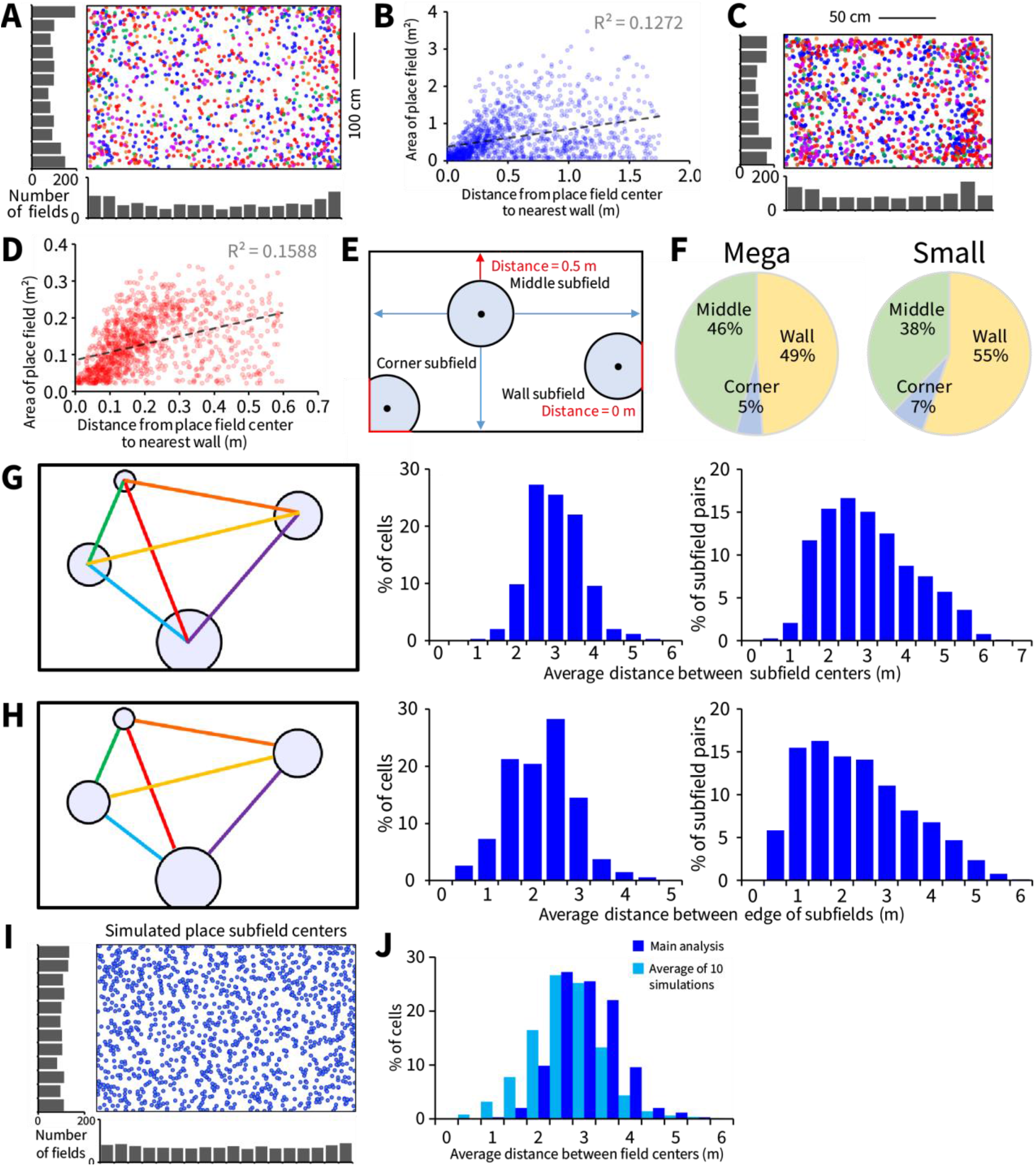
A well-isolated sample had comparable place cell characteristics to the population of cells and the distances between place cell subfields were normally distributed. (**A**) Plot of all place field centers in the megaspace, colors indicate place fields recorded from 5 different rats (blue, orange, red, green, purple). (**B**) Distance to the nearest wall plotted against subfield area for all subfields in the megaspace. (**C**) Plot of all subfield centers in the small environment. (**D**) Distance to the nearest wall plotted against subfield area for all subfields in the small environment. (**E**) The distance in each cardinal direction from the edge of each subfield to the maze walls was calculated. The red arrow shows the closest wall. Place subfields that contacted two, one, or no walls were designated “Corner”, “Wall”, and “Middle” subfields, respectively. (**F**) The megaspace and small environments had similar proportions of types of subfields. For place cells with at least 2 subfields, (**G**) the distance from the center of each subfield to the center of every other subfields (i.e. field pairs) and (**H**) the distance from the edge of each subfield to the edge of every other subfields were calculated. The average distance between subfields per place cell for both of these measures was normally distributed, whereas the distribution of distances between subfield pairs for both of these measures was right skewed, meaning a larger proportion of field pairs were closer together relative to the cell-averaged data. (**I**) Place field positions from the experiment are shown next to (**J**) an example of randomly generated positions. The distribution of average distance between field centers per cell was the same shape as the experimental data, but shifted towards higher values.

### Place cell representations were more dynamic in the megaspace than in classic environments

We next investigated the stability of the spatial representation in the megaspace after environment changes. We recorded 125 place cells from additional sessions in which two of the rats experienced the megaspace (Mega 1), followed by the small environment, followed by the megaspace (Mega 2) again (Fig. 5A). We compared the stability of place fields between Mega 1 and Mega 2 visits with the stability of fields between Small 1 and Small 2 visits in the main experiment sessions (Small 1 - Mega – Small 2). This analysis included all place cells defined from the main experiment sessions (n = 539). An independent t-test showed that place cells were less stable between megaspace visits than between small environment visits (*t*_(920)_ = 8.33, *P* < 0.0001; Fig. 5B). However, both populations included place cells that changed in size, position, and firing rates between environmental visits. This may be related to the large shift in environmental scale between the small and megaspace environments.

**Fig. 5:**
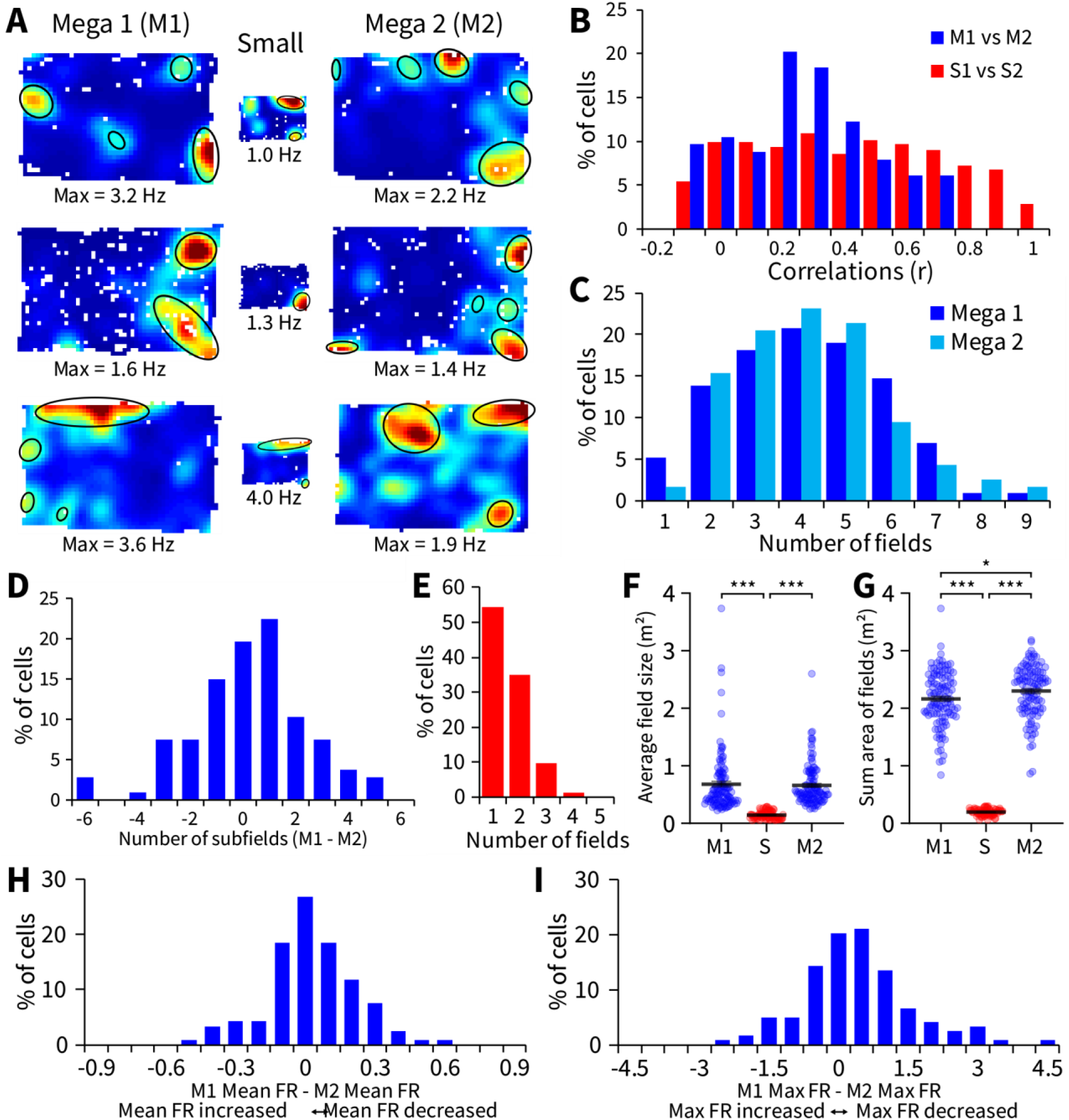
Place fields are more dynamic across visits in the megaspace than across visits in the small environment. (**A**) Representative examples of three place cells recorded in additional sessions in which rats foraged in the megaspace before (Mega 1) and after (Mega 2) the small environment. (**B**) Comparisons of rate map correlations, between the two megaspace visits (M1 vs M2) and the two small environment visits (S1 vs S2) from the main experiment. (**C)**The distribution of number of subfields per cell was comparable for the two megaspace visits. (**D**) Cell-to-cell variation in number of subfields (Mega 1 – Mega 2) between megaspace visits. (**E**) In the small environment, place cells had a similar distribution of number of subfields as in the main experiment. (**F**) The average size of subfields per cell was comparable between Mega 1 and Mega 2, however, (**G**) the sum area of subfields per cell was larger when the megaspace was revisited (Mega 2). Difference in (**H**) mean firing rate, and (**I**) maximum firing rate between megaspace visits for the population of cells. Negative values along the x-axis indicate increased firing in Mega 2 relative to Mega 1, whereas positive values indicate decreased firing in Mega 2 relative to Mega 1. All units in Hz. For all panels * = *P* <0.05, *** = *P* <0.0001.

We compared place cell characteristics between the three environmental visits using one-way Anova and found that the number of subfields did not vary between megaspace visits (*F*_(2,313)_ = 90.2, *P* < 0.0001; Tukey’s Mega 1 vs Mega 2, *P* = 0.37; Fig. 5C), and cell-to-cell variation in subfield numbers was unimodal but not normally distributed (Kolmogorov-Smirnov test; *D*_(114)_ = 0.13, *P* < 0.5; Fig. 5D). As expected, there were less subfields in the small environment (Tukey’s, Mega 1 and Mega 2 vs Small, *P* = 0.0001), which had a comparable distribution of subfield numbers as small environment visits in the main experiment (*F*_(1,847)_ = 0.79, *P* = 0.37; Fig. 5E and Fig. 2). The average area (*F*_(2,313)_ = 56.78, *P* < 0.0001; Fig. 5F) and sum area (*F*_(2,304)_ = 864.1, *P* < 0.0001; Fig. 5G) of subfields per cell were different between environment visits, which was driven by differences between the megaspace visits and small environment (Tukey’s, *P’s* < 0.0001). Although the average area of subfields per cell was comparable between Mega 1 and Mega 2 (*P =* 0.97), the sum area of place subfields was larger in Mega 2 than in Mega 1 (*P* < 0.05). However, the average (Fig. 5H) and maximum (Fig. 5I) firing rates were not different between megaspace visits (One-way Anova’s, Mega 1 vs Mega 2; Mean rate: *F*_(1,238)_ = 0.12, *P* = 0.73; Max rate: *F*_(1,238)_ = 1.22, *P* = 0.27). These findings suggest that both spatial and non-spatial associations may be more continuously updated [13] in large environments than in smaller ones. Some of these differences may also be associated with larger distances to anchoring cues in the megaspace, the different duration spent in the environments, or different time intervals between re-visits. These additional sessions also demonstrated that the multi-field place cell phenomenon was not specifically related to switching from small to subsequently larger environments, which has been the favored design in other studies [7, 9, 12].

### Place subfield properties are modulated by environmental scale

To study how the place cell representation changed with the scale of the environment, we recorded 130 additional place cells in two rats from sessions in which the environment size increased in three stages (Fig. 6A). In between navigating in the small environment and the megaspace, rats experienced a “large” environment which was intermediate in size (350 x 235 cm; 8.2 m^2^). As expected, the sum area of all place subfields increased as the environments expanded in size (One-way Anova; *F*_(2,336)_ = 1488.37, *P* < 0.0001; Fig. 6B). In contrast, the proportion of the environment covered by place fields per cell did not increase linearly (Fig. 6C); instead, it was similar for the two larger environments (*F*_(2,336)_ = 78.6, *P* < 0.0001; Tukey’s, Large vs Mega, *P* = 0.92). We found that the number of place subfields also increased with the scale of the environment (*F*_(2,387)_ = 133.22, *P* < 0.0001) with the highest proportion of cells exhibiting 1-2 subfields in the small environment, 3 subfields in the large environment, and 4-10 subfields in the megaspace (Fig. 6D). Variability in place field size also increased with environmental scale (Fig. 6E), whereas the ratio between the peak and average firing rate within place fields decreased slightly (*F*_(2,1128)_ = 9.97, *P* < 0.0001; 6F). The total area covered by a cells subfields was not correlated with the number of subfields and increased for larger environments (Fig. 6G). We compared the fraction of the environment covered by place fields for each cell in the small vs large (Fig. 6H), small vs mega (Fig. 6I), and large vs mega (Fig. 6J) environments, and found the correlations to be low, suggesting that they were unrelated. Across all recordings from the four different sized environments used in the study (Fig. 6K), the number of place subfields increased linearly with environment size (R^2^ = 0.9776; One-way Anova; *F*_(3,2152)_ = 608.8, *P* < 0.0001; all Tukey post-hoc comparisons, *P’s* < 0.001; Fig. 6L). Similarly, the sum area of all subfields per cell increased significantly (R^2^ = 0.9801; *F*_(3,2152)_ = 7774.6, *P* < 0.0001; all Tukey post-hoc comparisons, *P’s* < 0.0001; Fig. 6M) but with a strong exponential fit (r = 0.996) which matched the increase in area between the four environments (r = 0.987).

**Fig. 6:**
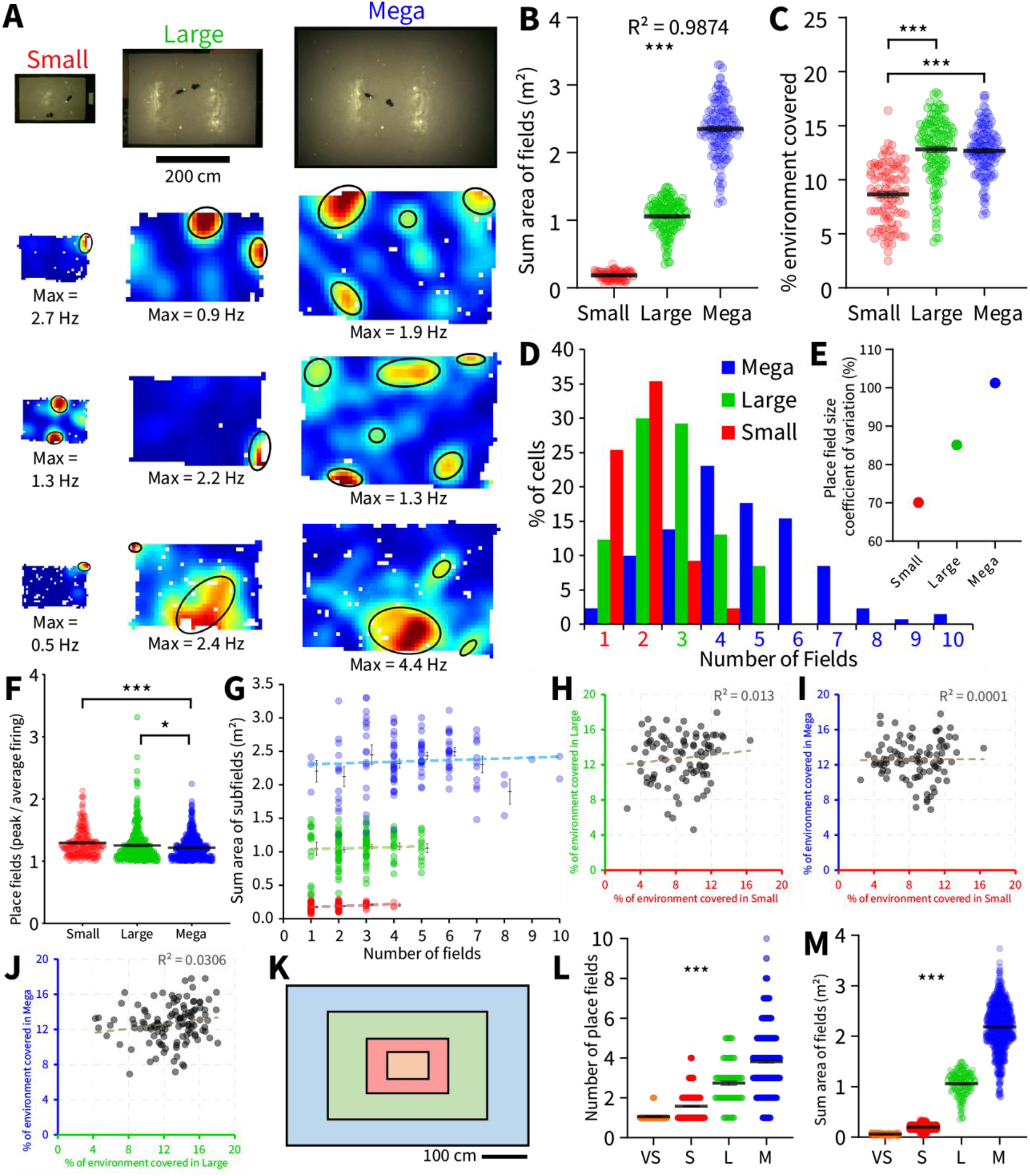
Place subfields scale with environment size. (**A**) Overhead view of three environments and example place cells from additional sessions in which the size of the environment increased in three stages. Between visiting the small environment and megaspace, a large environment (3.5 x 2.35 m) was visited that was intermediate in size. (**B**) The sum area of all subfields increased linearly as environment size expanded, however, (**C**) the percentage of the environment covered by subfields was comparable between the two larger environments. (**D**) Distribution of number of subfields per cell for the three environments. Number of subfields along the x-axis are color-coded to indicate which environment had the highest proportion of fields. (**E**) Variability of place field size increased with environment size. (**F**) The ratio of peak to average firing rate within place fields was comparable across environment sizes. (**G**) Distribution of sum subfield area for cells with different numbers of subfields in the megaspace (blue), large (green), and small (red) environments. The fraction of the environment covered by place fields was uncorrelated between the (**H**) Small and Large, (**I**) Small and Mega, and (**J**) Large and Mega environments. (**K**) To-scale depiction of the four environment sizes used in the current study, from smallest to largest: very small (VS, orange, 0.54m^2^), small (S, red, 2.16m^2^), large (L, green, 8.225m^2^), megaspace (M, blue, 18.55m^2^). Place cell recordings in these environments were aggregated from all session types (VS, n = 122; S, n = 1278; L, n = 130, M, n = 750). (**L**) As the environment size increased, the number of place subfields increased linearly and (**M**) sum area of subfields increased exponentially across the four environment sizes. For all panels * = *P* <0.05, *** = *P* <0.0001.

## Discussion

Our results show that the area of the environment covered by each dorsal CA1 place cell increases with the size of the environment, and that each cell is active in several distributed subfields of various sizes. The ability to exhibit different subfield sizes gives each place cell the capability to form a multi-scale representation of space. These multiple subfields also allow each cell to be active in several sections of the same environment, possibly spatially binding them, and allows for each location of the environment to be represented by a unique combination of subfields of different sizes [19]. Ensembles of dorsal CA1 place cells therefore form complex multi-scale codes capable of supporting concurrent and interdependent coarse- to fine-grained spatial representations, extending our current understanding of the hippocampal spatial code in large ethologically realistic environments.

The propensity for multiple place fields and up-scaling of field size increased as the environment size increased; an efficient way for a finite population of place cells to encode vast natural environments measured in kilometers [20, 21]. Place cells may be intrinsically multi-scale (multi-field) in all environments, even though only one or two place subfields can be physically reached by the animal in smaller ‘classic’ environments. An interesting question is how place cells with multiple spatial subfields can accurately represent the position of an animal? We found no within-field firing rate pattern that might explain how subfields from the same cell could be differentiated based on spiking activity. Instead, it is likely that overlaps from many different cell’s subfields use an ensemble pattern decoding scheme that can accurately estimate the animal’s current location [7, 12, 22]. Multiple subfields allow a cell to contribute to the ensemble in multiple regions of the environment at multiple scales. This raises the interesting possibility that in large environments, place cells may contribute to the spatial ‘binding’ of different subareas within the same environment, contributing to the animal understanding of the space as being the ‘same space’, whether it is physically located in one side of the room or in another [23]. The multiscale nature of the code also raises questions about the interactions of these CA1 cells with other types of cells known to be theoretically useful to spatial navigation, such as head direction cells, boundary vector cells [24] or landmark-vector cells [25].

Our work supports the findings of others showing that place cells exhibit multiple irregularly-arranged place fields in a large open environment [7, 12]. However, here we show a greater enlargement of place fields than would have been predicted and that in even larger environments, place subfields also greatly range in size, forming a representation at multiple spatial scales. Rich et al. showed multi-field place cells when rats traversed a winding 48m-long linear track [9], but did not report a multi-scale representation. Although the animals travelled a considerable linear distance when the track was fully extended, the total floorspace available to the animal was more than four times smaller than that of the megaspace. Furthermore, as cells were only recorded during one novel exposure session, direct comparison between the studies is difficult. It is likely that the encoding of novel environments is significantly different to that of a familiar one, at least in the requirement for the latter to retrieve and process memories. Rich et al. concluded, similarly to previous studies, that dorsal multi-field place cells may operate alongside a dedicated ventral hippocampal place cell population in order to encode differently sized environments.

Alongside others, we have also previously suggested a multi-scale representation of large-scale space involving the longitudinal axis of the hippocampus in which fine- and coarse-grained representations are supplied by the dorsal and ventral hippocampus, respectively [21]. However, considering the structural, connective, and functional gradients present along the dorsoventral hippocampus [26], it is likely that representations of different scales are in fact integrated along the entire hippocampus. Our results suggest this is indeed the case within the dorsal CA1. In correspondence to the manner in which single-field place cells increase in size along the dorsal-to-ventral hippocampal axis [1], we predict that the total area of the environment covered by multi-field place cells would also increase along the axis. Within large environments, the majority of ventral place subfields would be larger, but smaller place fields would also be exhibited, which could explain in part previous reports of smaller ventral place fields [27]. The concept of a dorsal-ventral functional gradient of small-to-large scale representations is challenged by our finding that individual multi-field cells within dorsal CA1 can exhibit a wide range of subfield sizes. Instead we propose that the multi scale coding is pervasive throughout the axis, and that place fields at all levels may be directly connected through the dense web of CA3 connections present along the longitudinal axis [21]. Large place fields at all levels may form distinct neural ensembles dedicated to encoding a lower-resolution and less computationally intensive representation supporting coarse travel. Simultaneously, longitudinal neural ensembles utilizing smaller place fields from these same cell populations are overlaid to provide higher resolution and details where needed within the environment [20]. Information selectively received by ventral levels (e.g. amygdala or prefrontal cortices) would then modulate all levels of the longitudinal axis simultaneously, at multiple scales. There is already evidence in human fMRI studies of fine- and coarse-grained hippocampal representations [28–30]. Interestingly, reliance on cognitive maps, and better navigational performance are related to greater posterior (dorsal), relative to anterior (ventral), hippocampal volume [31–33]. Other virtual navigation studies found that the anterior hippocampus became mainly involved when navigating through large and complex environments, whereas the posterior hippocampus was always active [28, 30]. Certainly, humans must make use of complex place cell maps utilizing three dimensions [34], over many overlapping spatial scales, from single rooms, to buildings, to streets, to cities, and beyond. It would be interesting to incorporate the concept of multi-scale place cells into models of how these hippocampal cells support networks of semantic cognitive space [35]. The idea of multi-scale overlapping place subfield ensembles may also be suited to understanding how mnemonic hierarchies may be encoded in autobiographical memory [16]. For example, memory of a life-event may constitute overlapping ensembles that encode both contextual (large subfields) and detailed (small subfields) features of the memory.

The increased navigational complexity inherent to the megaspace representation, which incorporates multiple subfields per cell and a wide range of subfield sizes, may require more flexibility and adaptive capability than previously thought when studying behavior in smaller environments. Our results suggest that place cell characteristics were more dynamic upon revisiting the megaspace compared to when revisiting the small environments, however this would need to be studied more directly, ideally with a second different megaspace room. The irregular patterns of place subfields observed in the current study suggests a flexible representation consisting of unique ensemble discharges of overlapping fields at any one location, rather than an orderly partitioning in which each region contains a field from each cell [7, 13].

Taken together, our findings reveal new coding properties and point to new ways in which place cells may operate in larger-scale navigational space and will require new generations of computational models of multiscale spatial navigation [13] and new experimental paradigms to be developed.

## Supporting information

VideoSummary

## Acknowledgements

This research is supported by: NSF CRCNS #1429937 and NSF IIS #1703340. We thank S. Gianelli and S. V. Srivathsa for technical assistance. We thank Dr. Bruce McNaughton and Dr. Andre Fenton for their constructive comments on previous versions of the manuscript.

## Author Contributions

B.H. and J.M.F. conceived and planned the experiments. B.H. performed the surgeries and collected the electrophysiological data with assistance from M.C. and M.S. B.H., J.M.F, and M.C. contributed to the interpretation of the results. B.H. led the analysis and writing the manuscript. All authors discussed the results and commented on the manuscript.

## Declaration of Interests

The authors declare no competing interests.

## Methods

All methods were approved by the University of Arizona IACUC and followed NIH guidelines.

### Subjects and behavioral apparatus

Five adult male Brown Norway Rats (6-7 months and 321-346 g at time of surgery) on a reverse 12/12 cycle were used in this study. Rats were trained to forage and follow a small robot (Fig. 1C) in a very large environment (530 x 350 cm). This ‘megaspace’ [36] was enclosed by black wooden walls (51 cm high). Large colorful national flags (71 x 56 cm) covered the east, west, and south room walls at varying heights, and irregular distances from each other. Smaller flags (~ 25 x 15 cm), cut into different shapes, were placed along all four maze walls at varying heights. All flags had different unique combinations of shapes and included light and dark colors. The floor of the room was painted with granular water-proof paint and contained multiple ‘cues’ in the form of small pieces of electrical tape of varying size and shape (~ 1–4 cm). See Fig. 1A for a top-down view of the megaspace. Flags and floor cues were chosen to provide a richly cued environment and were never displaced.

Three smaller environments were also used, consisting of modular walls centered within the megaspace, and sharing the same floorspace; these were designated as the ‘large’, ‘small’, and ‘very small’ environments. The large environment (350 x 235 cm) had 20 cm high wooden walls consisting of 3 segments per long side, and 2 segments per shorter side. Three different colors of segments were arranged so that the same color was never used for 3 adjacent segments, and that no corner or wall was the same. Some of the megaspace maze-wall flags, and all of the room-wall flags were visible from within the large environment. The small environment (180 x 120 cm) consisted of 33 cm high black wooden walls along three sides (north, east, and west) and a 51cm high black wall along the south side. A single white rectangular cue-card (21.6 cm high and 28 cm tall) was centered on the taller south wall. Only the larger flags positioned higher up on the room walls were visible to rats inside the small environment. The very small environment (90 x 60 cm) had black wooden walls 43 cm high with a white cue card (21.6cm high and 28cm tall) and an X painted in white paint opposite each other on the shorter walls. Maze and room wall flags were not visible from inside the very small environment.

The rat’s movements in the megaspace and large environment were captured by an overhead camera (PointGreyFlea3 at 25-30 frames per seconds) mounted on the ceiling in the center of the room. A separate overhead camera (Logitech Carl Zeiss Tessar Webcam HD 1080p, 25-30 fps) was used to capture the rat’s movement in the small and very small environments. The cameras provided inputs to our tracking software ZTracker, written in house in LabVIEW (National Instruments), and freely available from our website. A strip of LEDs near the cameras provided about 0.5-0.6 lux of light during the experiments.

### Sphero robot

The small robot used in the study was a Sphero 2.0 (Sphero, Boulder, CO) which was always fitted within a black plastic cart (Fig. 1C). A small black plastic weigh boat, containing mash (4:3 rat chow:water) was glued at the back of the cart. Sphero was linked via Bluetooth to custom in-house LabVIEW (National Instruments) software allowing the robot to be piloted with a joystick (Microsoft Sidewinder USB Joystick) enabling fine control of speed and trajectory. See [17] for more detailed information about Sphero, its control system, and integration with rat behavior. All control software to pilot the robot is available for download from our laboratory website. In the very small environment only, a smaller ‘mini-Sphero’ (Sphero, Boulder, CO; 4cm diameter) housed in a homemade 3D-printed cart was deployed to enable maneuvering in such restricted space (Fig. S4G). The homemade cart was 9.2 cm long, 5 cm high, and 5 cm at the widest point (the wheels). A small section of weigh boat was attached to the back of the cart creating a small dish in which mash (wet regular food) was placed, as with the regular-sized Sphero.

### Pre-Training

Rats were kept at 85-90 % of ad-libitum body weight and were fed after each training or recording session. Water was always available. After habituation to the environment in the home-cage for several days, rats were trained to sit on a towel-covered raised bucket lid (34.5 cm diameter, 83 cm high) in the center of the room for periods up to 1 hr. Next, as described previously [17], rats were trained to follow the Sphero robot while being habituated to the megaspace over several weeks. After two or three 10 – 15 min sessions following the robot in the megaspace and one session foraging in the small environment, the rats were put back on ad-libitum food in preparation for Hyperdrive implantation.

### General procedure

After surgical recovery (see below), rats were re-introduced to the various environments over the course of about a week. As the animals became more accustomed to the additional weight of the hyper-drive, small weights (9-32g) were slowly added to the drive’s protective cap to simulate the weight of the wireless headstage and build up the neck muscles. Elastic support, attached to the wireless headstage, was also used during training, mounted to the ceiling for the small environment, and attached to a long flexible pole held by an experimenter for the megaspace.

Each recording session began with a 10-20 min pre-rest period on the bucket, followed by three behavioural segments (visits to different environments; see ‘Session types’), followed by a 10-30 min post-rest period on the bucket. Within each session, the behavior in all three segments was either classical foraging or following the robot. In Sphero-following sessions, the robot was driven in front of the rat, maintaining a distance of ~ 15–25 cm, in a combination of straight and curving arcs around the environment (see Movie S1, and Fig. S1A-C and S3, for examples of the rat’s overhead path). The cumulative coverage of the room was monitored in real-time by the experimenter from the camera tracker. When the rat caught up with the robot it would slow or stop to allow the rat to consume food, if the robot was not caught, it would slow or stop after ~ 2-4 mins. When the weigh boat became empty, the robot was kept moving and interacting with the rat until the rat became unresponsive / disinterested or ~ 1 min had passed since the rat had fed, at which time the experimenter directed the robot to the edge of the maze, and re-baited the cart. In instances when the rat did not immediately follow the robot, simulated darting behavior [37] was used, eventually resulting in the rat following the robot. In classical foraging sessions, small 20 mg food pellets (TestDiet; Richmond, IN, USA) were tossed into the arena, and the rat was left to forage for the duration of each segment.

Cumulative tracking of the rat’s path was used to guide the animal to areas of the environment not covered sufficiently and influenced the length of each segment; longer segments were recorded if more coverage was needed. During the rest periods at the start and end of each session, the rat was placed on the bucket near the center of the room (center of both the small environment and megaspace). Between segments, the rat was placed on the bucket for 5-7 mins off to the side of the room while environments were erected / dismantled. The wireless head-stage was turned off during this time to allow it to cool down, and the battery was replaced if necessary. However, the headstage always remained connected for the duration of each daily session.

### Session types

In the main experimental sessions (Small 1-Mega-Small 2: S-M-S; n = 54 sessions), rats visited the small environment (8 – 10 mins), followed by the megaspace (35 – 55 mins), followed by the small environment again. These sessions compared place cell firing properties in the small and megaspace environments. In two of the rats, additional session-types were run. In eight sessions (7 Sphero-following, and 1 foraging), rats visited the megaspace, followed by the small environment, followed by the megaspace again (M-S-M). These sessions investigated the stability of place cell firing in the megaspace over several visits during the same session (Fig. 5A). In eight sessions (7 Sphero-following, and 1 foraging), rats visited the small environment, followed by the large environment for 25 mins, followed by the megaspace (S-L-M). These sessions investigated changes in place cell characteristics over three environments of increasing scales (Fig. 6A). In ten Sphero-following sessions, rats visited the small environment for all three behavioral segments (S-S-S). These sessions were used as control sessions for comparison with correlations performed between the small and megaspace revisits in other sessions (Fig. S4B). In three Sphero-following sessions, rats visited the very small environment (5 – 6 mins), followed by the small environment, followed by the very small environment again (V-S-V). These sessions established place cell characteristics in a constrained environment, traditionally used for recording place cells (< 1m^2^; Fig. S4F).

### Surgery and recording techniques

After completion of pre-training, rats were anesthetized using 2–3% isoflurane in oxygen, placed in a stereotaxic frame, and implanted with a Hyperdrive [17, 38] aimed at the right dorsal CA1 hippocampal cell body layer (−4.75 mm posterior, 4.0 mm lateral to bregma, 10° angle away from midline). The drive was anchored to the skull with seven anchor screws and dental acrylic, and two of these screws were used as animal grounds. Additionally, two EEG electrodes (Teflon-insulated stainless-steel wire, 0.0045 in.) were implanted in the right medial prefrontal cortex (+3.00mm posterior, 1.2 mm lateral to bregma, 2.8 mm depth, 9° angle towards midline). An EMG electrode was implanted in the neck muscles of the rat to help assess sleep during the rest phases (data not shown). All implantation coordinates were modified proportionally to the Bregma-to-Lamda distance of the animal using a brain atlas [39]. Glycopyrolate (I.M.) was administered during the surgery to alleviate congestion, and Carprofen, an analgesic, was given (I.P.) during surgery and again the day after.

The Hyperdrive contained 14 independently movable tetrodes, two of which were used as reference. Tetrodes were constructed from four strands of insulated wire (12 μm diameter nickel-chrome wire), gold-plated to reduce wire impedance to 0.5 MΩ (at 1 kHz). Following surgery, about 4-6 tetrodes at a time were slowly lowered in batches toward the hippocampal dorsal CA1 pyramidal cell body layer both to facilitate recordings over several months and to avoid instability. Reference tetrodes were left in an electrically quiet zone in the cortex or corpus callosum. Tetrodes were spaced ~50μm apart, and lowered at the end of each experimental session, to ensure that the same cells were not recorded in multiple sessions.

Electrophysiological recordings were made using either a wireless Cube 64 or Cube 2 headstage (currently renamed ‘Freelynx’, Fig. 1C shows the Cube 2 headstage mounted on the hyperdrive of a moving rat) controlled by a Digital Lynx SX system (Neuralynx, Bozeman, MT). Single-unit data was amplified, filtered (600–8000 Hz), and digitized at a rate of 30 kHz. Local field potential was recorded from one channel per tetrode, filtered between 0.5 – 450 Hz, digitized at 2 kHz, and used to detect the presence of sharp wave ripple oscillations, confirming that tetrodes were in the dorsal CA1 cell body layer. Two LEDs (red/green) mounted on the headstage were used to track the animal’s movements with the overhead cameras.

### Spike sorting

Action potentials were sorted offline using Spike2 software (CED, Cambridge UK) and further analyzed using custom Matlab code. Clustering was performed manually by a single experimenter in three-dimensional projections based on the principal components of the waveform amplitude. Data from each session – the three behavioral segments and two rest periods – were spike-sorted together. Only well isolated clusters with pyramidal waveforms, signal-to-noise ratio of at least 4 on one of the 4 channels were retained. Signal was measured as the mean amplitude of the action potential (peak-to-trough), and noise was measured as the mean amplitude of the initial 2 points of each waveform. Clusters isolated from the same tetrode were manually checked to insure each had a sufficiently different configuration of shape/amplitudes across the four channels. Clusters were labelled as either putative excitatory cells or putative interneurons using differences in spike width, average firing rate and complex-spike bursting.

### Detecting behavioral stops and sharp wave ripples (SWR)

Position data, based on tracking of the LEDs on the head stage, were analyzed and all stop periods were detected. Stops were designated as periods when instantaneous velocity dropped below 6 cm/sec for a period of at least 0.5 sec. SWR events were detected using the best two LFP channels per session which were band pass filtered between 100-250 Hz and SWR envelopes calculated using a Hilbert transform, smoothed with a Gaussian kernel (3ms standard deviation). During behavioral segments, SWR events were detected as times within stop periods when the smoothed envelope exceeded 4 standard deviations above the mean for at least 20 ms. For rest segments, SWR events were smoothed envelopes exceeding 2 standard deviations above the mean for at least 20 ms during stop periods only. SWR events included 10 ms before and after the envelope, and envelopes exceeding 11 standard deviations above the mean were rejected as artifacts. All spikes occurring during sharp wave ripples were removed when generating spatial-firing rate maps to avoid any SWR activity contamination [40].

### Ratemaps and place fields

The position data for each session was sorted into bins of 12 x 12 camera pixels (5 x 5 cm for the small and very small environments, 10 x 10 cm bins for the megaspace and large) with a velocity threshold of 10 cm /sec [41]. Spike-count and occupancy maps were computed for each cell by counting the number of spikes occurring in each spatial bin, and the time spent in each spatial bin, respectively. Spike-count bins containing only one spike and occupancy bins visited for less than 0.08 secs, were considered empty. Both maps were smoothed using a square Hanning kernel window and the final place field map was produced by dividing the smoothed spike-count by the smoothed occupancy. The peak firing bin for each cell was used to colour code the spatial-firing rate map from dark red (highest firing) to dark blue (lowest firing).

The spatial information content (bits/spike) of spatial-firing ratemaps was calculated [42]. The occupancy map was used to quantify the spatial coverage (% Occupied bins) quality of each behavioural segment in each session by calculating the percentage of filled occupancy bins.

Cells were classified as ‘place cells’ only if: (i) mean firing rate was >0.1 Hz but <5 Hz, (ii) spatial information content >0.5 in at least one recorded environment [43, 44], (iii) they possessed pyramidal waveforms, which were manually checked in all cells, with (iv) signal-to-noise >4 on at least one tetrode channel.

Place fields were then designated as disconnected rate map regions of high activity > 200 cm^2^, with firing rate threshold >1.2 standard deviations above the mean firing rate in all bins using the *regionprops()* function in Matlab (Mathworks). The centroid pixel coordinates (x,y), and area (cm^2^) of this region were used to plot an ellipsoid fitted around the edges of each field to aid with visualisation of the place fields. The highest firing rate bin was designated as the maximum firing rate for each subfield. For each place cell with at least 2 place subfields, the absolute difference in maximum firing rate between each possible pair of subfields was determined (Fig. 3I).

For the S-M-S and S-S-S sessions, only place cells with correlated firing-rate maps between the two smaller environments were retained for analysis. Pearson correlations were calculated between the small environment rate maps recorded before (Small 1) and after (Small 2) exposure to the megaspace. This correlation was used to calculate a z-score by comparing it to correlations generated from 300 shuffled versions of each rate map in which the bins were spatially shuffled randomly. Eligible place cells had to have a z-score greater than 2.5, placing them above the 99.5% percentile cutoff of the shuffled distribution. For the other session types (M-S-M, S-L-M, V-S-V), cells that were not active (<0.1 Hz) in some of the environments but were otherwise eligible as place cells, were included in the analyses.

Distance between pairs of fields (in the same environment) was calculated both as the Euclidean distance between the centroids of each field, as well as the distance between the edges of each field, by subtracting the radius of each ‘idealised’ (circular) field from the first measure (Fig. 4G, H). Similarly, distance of each field to the closest wall was the shortest straight-line cardinal distance (x and y) from the centroid to each of the four walls, with and without the addition of the field’s radius. Using these measurements, fields were designated as ‘wall fields’ if the fields edge contacted the wall (distance =< radius) in one direction, ‘corner fields’ if contacting the wall in two directions, and ‘middle fields’ if they did not contact the wall (Fig. 4E).

### Rate map correlations and shuffles

Additional correlations were computed for each type of session to compare rate maps in the different sized environments. In each session, tracking data from all behavioral segments were re-sized to the same dimensions of data recorded in the smallest environment employed during that session (Fig. S4A). Resized rate maps were generated in the same way as the small environment rate maps and then compared using Pearson correlations and z-score comparisons against shuffled maps (300 shuffles).

In the main experimental sessions (S-M-S), comparisons were also made between small environment rate maps and the cell activity in the larger environments restricted to the same floor space only (Fig. S4D). This was achieved by re-scaling the megaspace tracking data to the pixel/cm scale of the small environment (0.46 cm / pixel) and generating new cropped rate maps encompassing cell spiking and occupancy only in the megaspace floor space occupied by the small environment (shown by yellow dotted line in Fig. 1A). These were compared with Small 1 and Small 2 ratemaps via Pearson correlations and z-score comparisons against shuffled maps (300 shuffles).

### Well-isolated place cell population

The well-isolated population subsample of 125 place cells from the main analysis included only 1 cell from each active tetrode per session (isolated cells with highest signal-to-noise ratio). This was done to eliminate any potential spike-cutting error. The sample population included contributions of cells from Sphero and foraging sessions, and from each animal, that matched the proportion of cells contributed by each to the total population of 383 cells, except for one rat that had only 2 foraging sessions with high cell yields, which contributed 3 additional Sphero sessions instead of foraging sessions. This well-isolated sub-population was compared to the main analysis population to ensure that findings in the megaspace were not due to multiple cells being clustered together.

The well-isolated subsample was also used to visualize a population of place fields in the three environments by plotting each place field’s center and area as semi-transparent ‘idealized’ circles of the same area as each place field (Fig. 3A). The 532 place subfields exhibited in the megaspace were split into seven even ranges based on their area, which were color-coded from purple for smallest to red for largest. These color-coded size ranges were then applied to the 219 subfields in Small 1 and the 209 subfields in Small 2. The area ranges for the color coding was: Purple <0.092m^2^; Dark Blue: <0.21m^2^; Light Blue <0.366m^2^; Green <0.54m^2^; Yellow <0.81m^2^; Orange <1.22m^2^; Red <3.47m^2^. When the entire population was plotted, it became graphically difficult to distinguish individual fields, however the field centers from all cells are shown for the megaspace (Fig. 4A) and small environment (Fig. 4C) color-coded by animal.

### Ensemble place field overlapping

We plotted the well-isolated subsample population of subfields from the main experiment as borderless circular fields with an alpha level of 0.05 in order to quantify the amount of overlaps between place fields (Fig. 3C). This provided a measure similar to % of environment covered by place fields, but also took into account the density of place fields at every pixel location throughout the different environments. The image was inverted and pixel density was analysed using Image J (NIH). To help identify the pixel intensities relating to specific number of subfield overlaps, a test figure was generated in which 60 overlapping place subfields with the same alpha level, of diminishing size, were plotted at the same location. Analysis of the test figure produced 60 peak intensities corresponding to the levels of overlap ranging from intensity values of 13, for one overlap, to 245 for 60 overlaps, along the 255-pixel intensity scale. Pixel intensity counts from the data were binned evenly around these peak values for small environment and megaspace subfield plots, which included peak intensities that matched the test figure. For the subsample population, the distribution of overlaps in Small 1 and Small 2 was comparable (*F*_(1,197)_ = 1.31, *P* = 0.72), so were averaged and compared directly to the megaspace overlaps.

### Histology and tetrode placement

The correct position of the electrode tips were confirmed in all animals by small electrolytic lesions on each of the tetrode wires (30 μA, 8-s positive to electrode, negative to ground) both the day before and just prior to the perfusion. Animals were then deeply anesthetized with a Ketamine/Xylazine mixture (0.45 and 0.05 mg/kg respectively) and transcardially perfused through the left ventricle with a Heparin-saline flush (200 ml) followed by 250 ml of cold 4% paraformaldehyde in 0.1M phosphate buffer (pH 7.4). After the brain was removed, it was post-fixed in the same fixative for 1 day and then transferred to a solution of 30% sucrose in PBS (phosphate buffer 0.01 M, NaCl 0.9%) with 0.02% sodium azide. At a later date, brains were then blocked in the coronal plane and immediately cut with a Cryostat (Leica) set for a thickness of 30-50 μm. Every section was obtained from the region of the EEG electrode track in the medial prefrontal cortex (data not shown), and the region encompassing the hyperdrive bundle in the hippocampus, and stained with cresyl violet (Nissl) then mounted on slides and cover-slipped [38].

Each tetrodes intersection with the hippocampal dorsal CA1 was recorded on digital photomicrographs (Stereo Microscope, 10x magnification) by comparing tetrode traces and electrolytic lesions on successive sections (Fig. S2 shows tetrode positions in dorsal CA1 for all rats). Each set of coronal photomicrographs was compared to brain atlas plates [39] to estimate the anterior / posterior position within the dorsal hippocampus.

### Data Analysis

Analysis of place field characteristics between environments and comparison of cells recorded during robot-following and foraging sessions were done using ANOVA with an alpha level of *P* < 0.05. Tukey’s post-hoc tests were used to test for group differences, where applicable. Kolmogorov-Smironov tests were used to test normality of frequency distributions. Correlation coefficient (r) and coefficient of determination (r^2^) were used to measure the statistical relationship between variables and to determine best fits. Comparison of the degree of subfield overlap between environments used an ANOVA in which each level of overlap was weighted by the fraction of environment covered. Distributions of place cell ratemap correlations between environment re-visits (i.e. Small 1 vs Small 2 or Mega 1 vs Mega 2) were compared using independent t-tests. All statistical test were performed in SPSS.

---

**Fig. S1:**
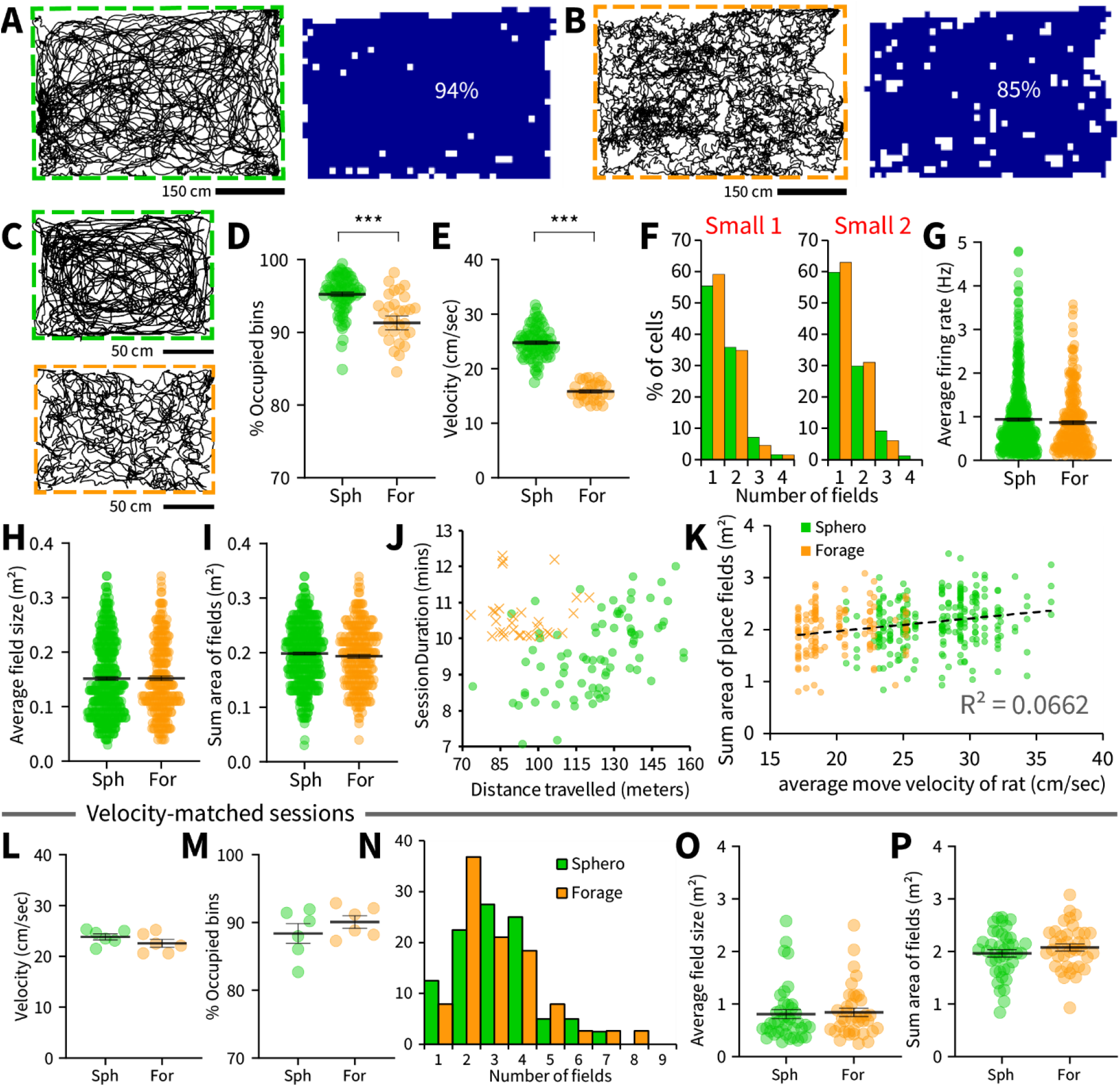
Robot-following facilitated greater speed and coverage in both environments. (**A**) Examples of tracking data from a representative ‘Sphero’ robot-following and (**B**) a representative foraging session in the megaspace. On the right of each, the occupancy map for each session that were used to calculate “coverage” of the environment. Percentages show the ratio of filled (blue squares) compared to empty (white squares) occupancy bins. (**C**) Example trajectories in the small environment from a (top) robot following, and (bottom) foraging session. (**D**) The fraction of the room covered by the occupancy map and (**E**) rat average velocity in the small environments, were greater during robot-following than during foraging sessions (One-way Anova’s: *F*_(1,53)_ = 54.26, *P* <0.0001 and *F*_(1,53)_ = 25.43, *P* <0.0001, respectively). (**F-I**) However, there was no significant difference in place cell characteristics in the small environment between robot following and foraging sessions (number of fields, *F*_(1,764)_ = 2.13, *P* = 0.15; average firing rate, *F*_(1,764)_ = 1.48, *P* = 0.22; mean size of place fields, *F*_(1,764)_ = 0.001, *P* = 0.97: sum area of place fields, *F*_(1,764)_ = 1.69, *P* = 0.19). (**J**) Session duration (mins) vs distance travelled (m) is plotted for small environment recordings; Sphero-following shown in green circles, foraging shown in orange crosses. (**K**) The sum area of subfields in the megaspace for all robot following and foraging sessions is plotted against the velocity of the animal. The weak positive correlation (r = 0.26) suggested that the lower sum area of fields in the megaspace for foraging sessions may be related to lower average velocity. Therefore, we compared a sub-set of velocity-matched robot following and foraging sessions (n = 6 each), in which (**L**) velocity and (**M**) fraction of occupied bins were not significantly different from each other (One way Anova’s, Average velocity: *F*_(1,10)_ = 1.69, *P* = 0.22; % Occupied bins: *F*_(1,10)_ = 0.97, *P* = 0.35). (**N**) The number of subfields (*F*_(1,76)_ = 0.0004, *P* = 0.98), (**O**) average subfield size (*F*_(1,76)_ = 0.08, *P* = 0.77), and (**P**) sum area of subfields (*F*_(1,76)_ = 1.34, *P* = 0.25) per cell did not differ for the 40 robot following place cells and 38 foraging place cells from these velocity-matched sessions. For all panels ** = *P* <0.001, *** = *P* <0.0001.

**Fig. S2:**
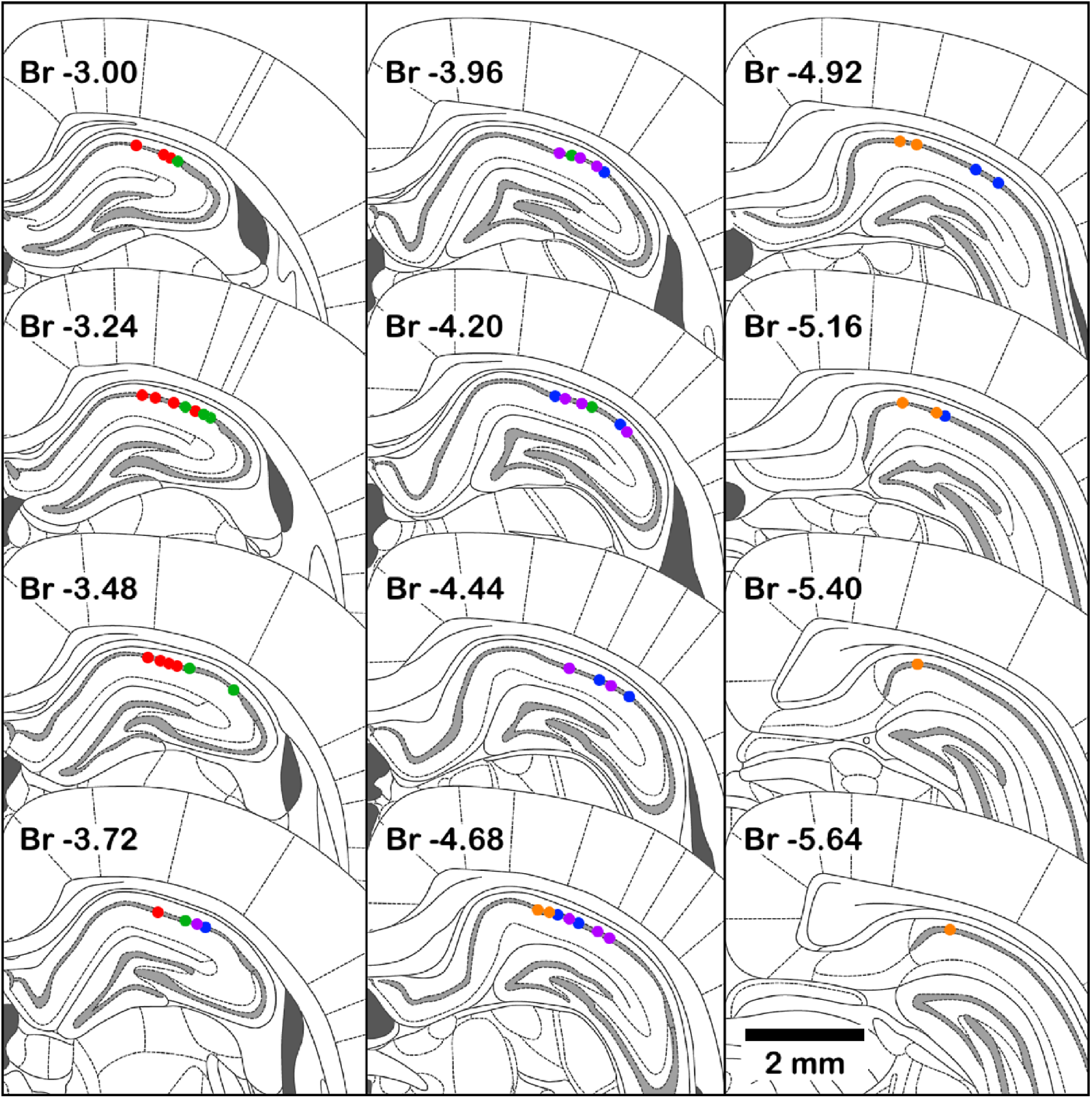
Tetrode locations in the dorsal CA1 cell body layer. Coronal atlas plates (Paxinos and Watson, 2006) are shown, ordered moving down through the columns from left to right in steps of 0.24 mm, with the distance from bregma (mm) shown for each plate. Tetrode intersections with the CA1 cell body layer were marked on photomicrographs of cresyl violet stained sections taken from each rat which were matched to all available Atlas Plates staggered in distances of 0.12 - 0.16 mm apart (more plates than are shown here). All tetrode positions in dorsal CA1 are shown on the closest template in five colors, one for each rat. Three tetrodes were found to be located outside CA1, in the CA2 region, and were not included in the data set (not shown).

**Fig. S3:**
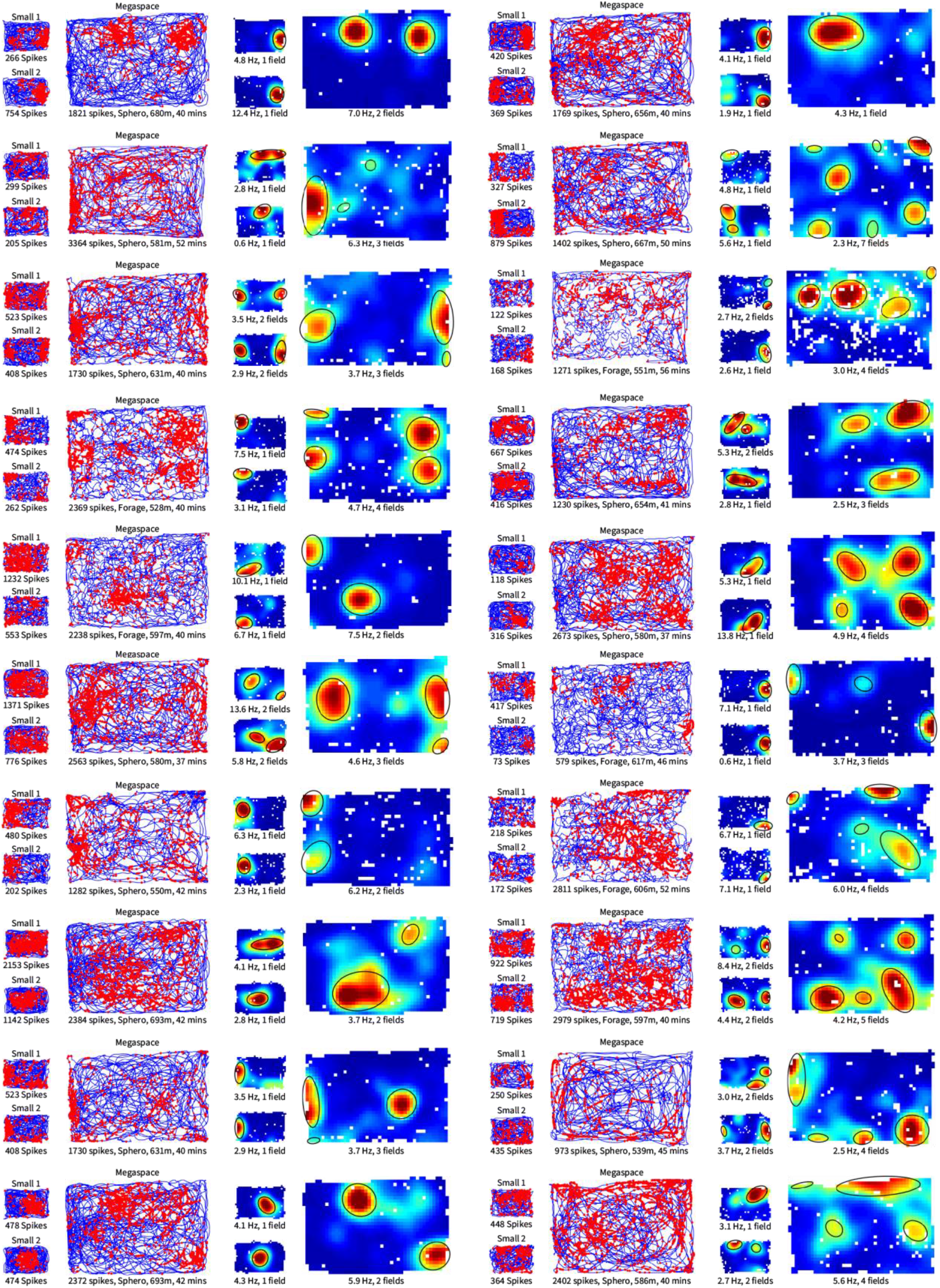
Additional example place cells with raw spike plots. These example cells show a representative sample of the variety of place cell firing observed both in the megaspace and small environments. Twenty different place cells are shown from the main experimental sessions (Small 1 – Megaspace – Small 2, for each cell) in 2 columns of 10 rows. On the left side of each example, the trajectory of the animal is shown (blue line) with cell firing plotted on top (red dots). On the right side of each example, the firing rate maps are shown; the peak firing rate (Hz) and number of place subfields are listed underneath each. High firing rates are represented by hot colours, and white bins show regions that were not covered sufficiently by the animal. Ellipsoids are plotted based on the parameters of each place field to aid with visualization.

**Fig. S4:**
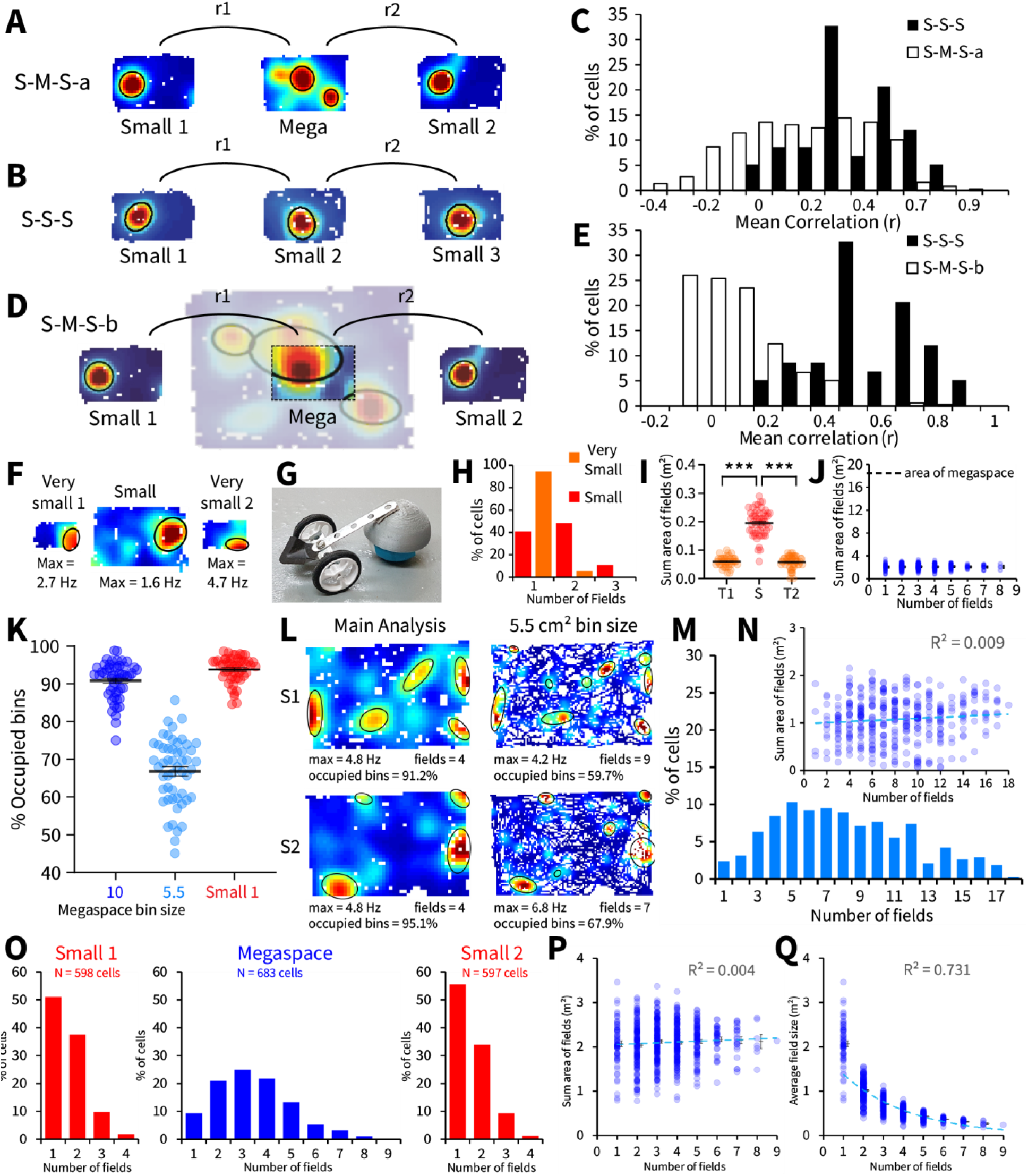
Place cell firing in the megaspace is unrelated to the small environment, and cannot be explained by the recording setup, bin-size, or criterion for cell inclusion. (**A**) We examined whether place cell firing in the different environments was related by re-scaling the megaspace place maps and directly comparing with the same cell’s spatial firing in the small environments. Pearson correlations between Small 1 and the re-sized megaspace map (r1), and Small 2 and the re-sized megaspace map (r2) were averaged for each cell (S-M-S-a). (**B**) These correlations were compared to r1 and r2 averaged for each place cell from control sessions in which the small environment was experienced three consecutive times (S-S-S). (**C**) The distributions of S-S-S and S-M-S-a correlations were significantly different (*t*_(920)_ = −11.45, *P* < 0.0001). (**D**) We also looked at whether place cell firings in the small environments were related to their firing in the specific section of megaspace floor-space shared by the small environments (S-M-S-b; yellow-dotted lines in Fig. 1A). (**E**) Again, we found that distributions of S-S-S and S-M-S-b correlations were also significantly different (*t*_(920)_ = 18.38, *P* < 0.0001). (**F**) Example place cells recorded in additional sessions in a very small environment (90 x 60 cm), before and after the small environment (180 x 120 cm). (**G**) In the limited space of the very small environment, a smaller version of the robot (Mini Sphero) housed inside a homemade cart was used. (**H**) Similarly to previous studies using a ‘classic’ <ce cells had more subfields in the small compared to both very small environments which had comparable numbers of fields (*F*_(2,141)_ = 35.75, *P* < 0.0001; Tukey post-hoc tests, Small vs. Very small 1 and Small vs. Very small 2, *P’s* <0.0001, Very small 1 vs. Very small 2, *P* = 0.98). (**I**) Total area of all place fields was greater in the Small compared to both Very small environments which did not differ (*F*_(2,141)_ = 298.05, *P* < 0.0001; Tukey post-hoc tests, Small vs Very small 1 and Small vs Very small 2, *P’s* < 0.0001, Very small 1 vs Very small 2, *P* = 0.96). (**J**) The same graph from Fig. 2I showing the sum area of place fields for cells with different numbers of subfields, but with y-axis extended up to the size of the megaspace (18.55 m^2^). (**K**) We used a larger bin size for the megaspace compared to the small environment in order to normalize the occupancy between the two environments. Using the lower bin size (5.5) in the megaspace resulted in greatly reduced % coverage of the space to levels traditionally not acceptable for place field computation (<90%). Using such small bin sizes would require running the animal much longer, to extents not possible with the current wireless technology. We re-analyzed the megaspace, using the same bin-size as we used in the small environment. (**L**) Two example cells are shown with the larger bin size on the left, and smaller bin size on the right. The smaller bin size had more place subfields, some of which appear to be ‘true subfields’ correctly separated at the lower bin size, whereas others are incorrectly separated due to more gaps in the place map from unoccupied bins. (**M**) Despite the greater number of subfields per cell in the megaspace, (**N**) the sum area of subfields per cell and numbers of subfields per cell remained uncorrelated. In our main analysis, we excluded place cells that did not fire during all three environment visits, as well as place cells with insufficiently stable firing between Small 1 and Small 2. Here, we re-analyzed the data to include this larger cell population in order to verify that our findings are not associated with the place cell criterion used in the manuscript. (**O**) This larger cell population had a comparable distribution of place fields to the more restricted cell population for the megaspace and Small 2, but the Small 1 distribution differed (One way Anova’s; Megaspace: *F*_(1,1064)_ = 0.53, *P* = 0.47; Small 2: *F*_(1,978)_ = 2.54, *P* = 0.11; Small 1: *F*_(1,979)_ = 4.23, *P* < 0.05). The difference in the small environment for the larger cell population was a slight decrease in the proportion of cells with only 1 subfield, and a slight increase in the proportion of cells with 2-4 subfields. (**P**) The sum area of subfields and number of subfields per cell remained uncorrelated and (**Q**) the relationship between average place field size and number of place fields per cell stayed consistent with the larger cell population. For all panels *** = *P* <0.0001.

**Fig. S5:**
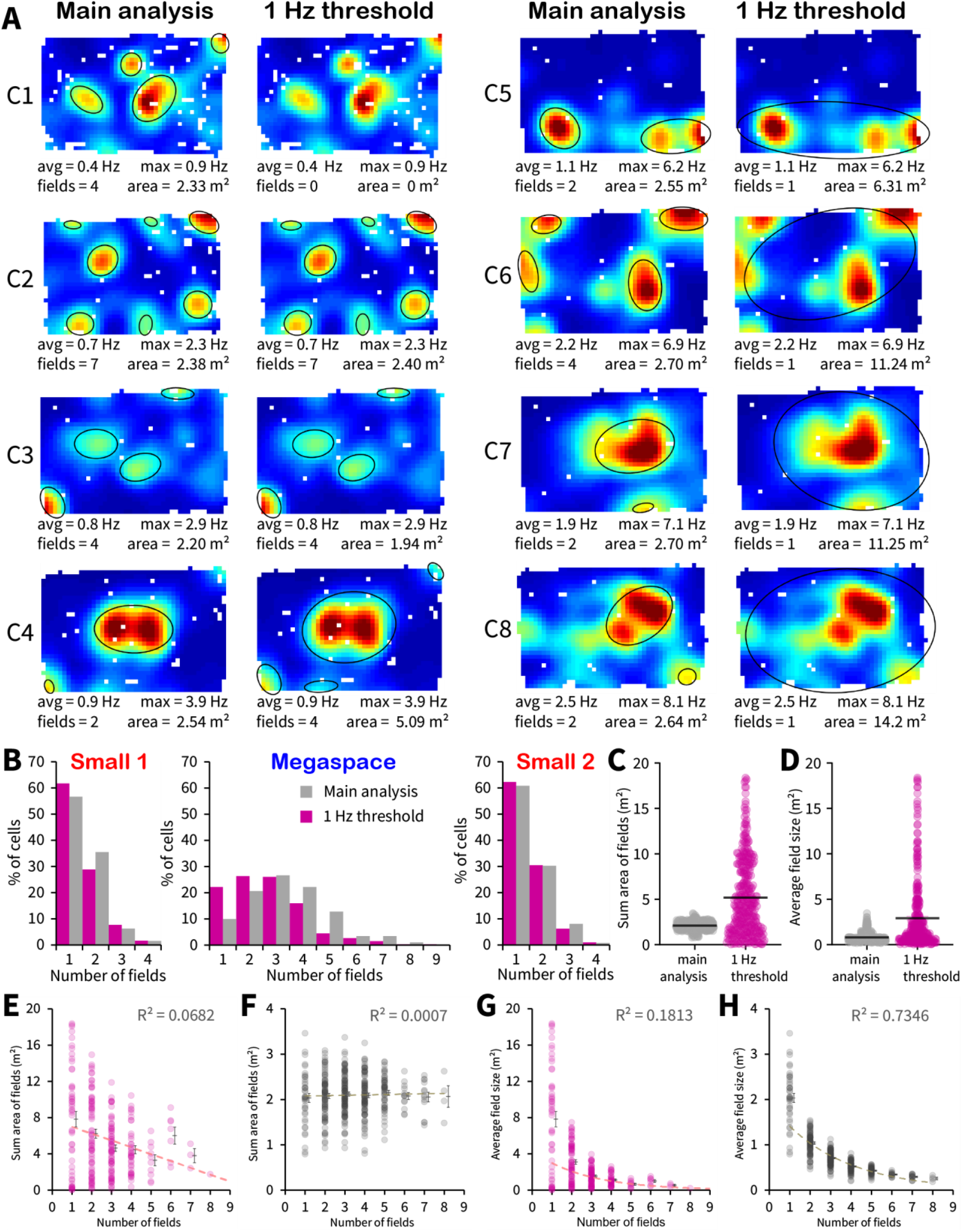
Place subfields still ranged in size, and cells with different numbers of subfields had similar average sum area when fields were defined with a fixed 1 Hz threshold. (**A**) Multiple example cells (C1 – C8) are shown showing place fields designated using the standard deviation method from the main analysis on the left side of each example, and place fields designated with a 1 Hz common threshold on the right side of each example [S1]. The examples cells are shown in order of firing rate from low (C1) through to high (C8). The common threshold method gives similar results to the mean + 1.2 standard deviation method for cells with intermediate values for maximum and average firing rates like C2 and C3. For cells with higher firing rates, the common threshold can label additional fields that fell below the threshold using the standard deviation method, see C4, but in many cases also combine fields that appear separate in the place map, such as with cell C5. This is because the permissive 1 Hz criterion props up weak bins in between strong bins, and this propping-up links the subfields together instead of separating them. This linkage produces abnormally large fields. Low firing rate place cells like C1 are not labelled with any place fields using the common threshold method; 45 out of 383 place cells in the megaspace (13.3%) were labelled with 0 place fields, and these ‘non-active’ cells are not included in the subsequent panels. For the highest firing rate place cells, the common threshold joined up place fields that appear separate on the place map to produce very large fields and sum areas as in C6, C7, and C8. (**B**) The 1 Hz common threshold resulted in a different distribution of number of fields in the megaspace with more cells having only 1 or 2 fields, but in the small environments the distribution of number of fields is comparable to place field labelling with the standard deviation method. There was a larger range in (**C**) sum area of subfields per cell and (**D**) average subfield size per cell in the megaspace using the fixed 1 Hz common threshold method, due to some very large place fields contributed by the higher firing rate cells. (**E**) There was a tendency for average sum area of place fields to reduce the more subfields a cell had compared with (**F**) the same graph produced using the cell-specific standard deviation method. (**G**) The tendency for average place field size to reduce, the more subfields a cell had in the megaspace, also persisted, but was weaker and noisier than when (**H**) the standard deviation method was used to label place fields in the megaspace.

**Fig. S6:**
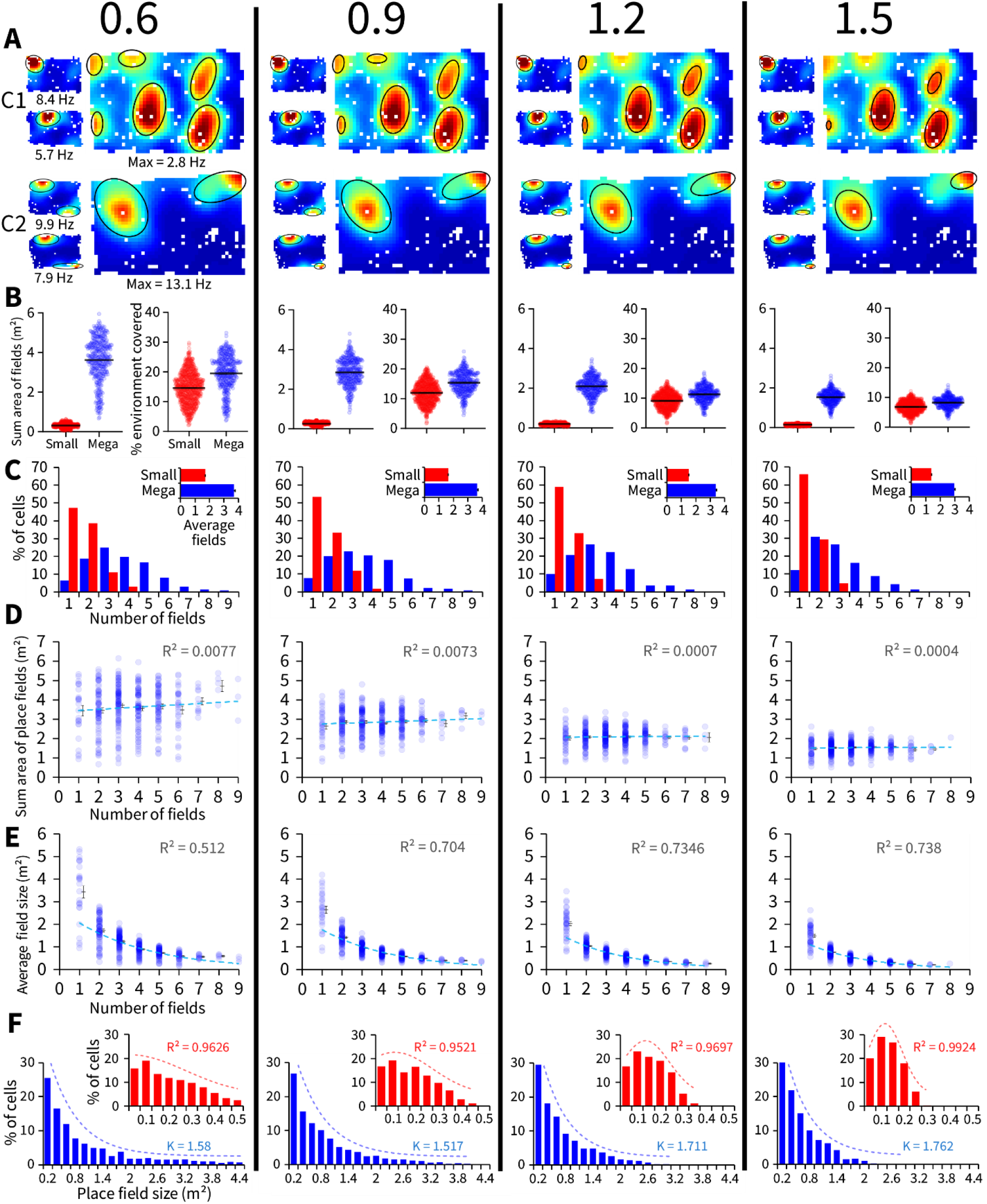
The differences between place subfield structure in the small environments and megaspace were consistent across different thresholds for place field labelling. (**A**) Examples of place maps from two place cells (C1 and C2) thresholded at 0.6, 0.9, 1.2, 1.5 (left to right columns) standard deviations above their mean firing rate for the identification of place fields. Small 1 is shown in the top left, and Small 2 is shown in the bottom right for each panel. Cell C1 gives an example of how different numbers of fields can be labelled as the threshold changes. Cell C2 gives an example of a cell in which only the size of the subfields change with threshold, but the number of subfields remains robustly constant in both small and large environments. (**B**) Sum area of all place fields and percentage of the environment covered by place fields are shown for each cell in the small environment (red), and megaspace (blue). (**C**) Number of place subfields per cell and average for all cells are shown in inset. As expected, the higher the threshold used, the smaller the area of subfields, and the fewer the number of subfields identified in both the small environment and megaspace. These effects were more pronounced in the megaspace due to its higher ceiling for these values. Across the different thresholds, (**D**) the sum area of all subfields and the number of subfields a cell had in the megaspace remained uncorrelated. (**E**) Similarly, the gradual decay of average field size in the megaspace as a function of the number of subfields persists. (**F**) Distribution of subfield sizes for the population of subfields pooled from all cells in the megaspace (blue) and small (red, inset) environments. In the megaspace this distribution was well fitted by a negative exponential curve, the rate of decay (K =) is listed for each threshold. In the small environment the distribution was gaussian at higher thresholds, and more linear at lower thresholds, the coefficient of determination (R^2^) is listed for each threshold.

## Movie S1: Title and Short Legend

**Movie Title:** Place Cell Recordings in a Megaspace

**Short Legend:** Place cells are wirelessly recorded from the hippocampus in rats as they follow a small robot. Between small environments, the rat is recorded in a very large ‘megaspace’. Place cells in the megaspace cover a similar total area but are fragmented into different numbers of subfields. These subfields vary in size, so that the population of place cells forms a multi-scale representation in the megaspace capable of supporting both coarse-and fine-grained representations of the environment. Additional recordings in environments of increasing scales show that the total area covered by each place cell is comparable within each environmental scale.

## Notes

http://amygdala.psychdept.arizona.edu/DataDemos.html

### Competing Interest Statement

The authors have declared no competing interest.

### Summary of Updates

Supplemental materials and movie

